# Unbalanced response to growth variations reshapes the cell fate decision landscape

**DOI:** 10.1101/2022.09.13.507864

**Authors:** Jingwen Zhu, Pan Chu, Xiongfei Fu

**Affiliations:** CAS Key Laboratory for Quantitative Engineering Biology, Shenzhen Institute of Synthetic Biology, Shenzhen Institutes of Advanced Technology, Chinese Academy of Sciences, Shenzhen 518055, China; University of Chinese Academy of Sciences, Beijing 100049, China

**Author notes:** These authors contributed equally: Jingwen Zhu, Pan Chu.

## Abstract

The global regulation of cell growth rate on gene expression perturbs the performance of gene networks, which would impose complex variations on the cell-fate decision landscape. Here, we utilize a simple synthetic circuit of mutual repression that allows a bistable landscape, to examine how such global regulation would affect the stability of phenotypic landscape and the accompanying dynamics of cell-fate determination. We show that the landscape experiences a growth-rate-induced bifurcation between monostability and bistability. Theoretical and experimental analyses reveal that this bifurcating deformation of landscape arises from the unbalanced response of gene expression to growth variations. The path of growth transition across the bifurcation would reshape cell-fate decisions. These results demonstrate the importance of growth regulation on cell-fate determination processes, regardless of specific molecular signalling or regulation.

## Introduction

Cell fate determination is a ubiquitous and important process in living systems, enabling cells to transit to different, functionally important, and heritable states in the absence of genetic variations^1,2^. The classical theory proposed by Waddington^3^depicts the developmental process as a ball rolling down the epigenetic landscape with ‘hills’ and ‘valleys’, of which the ‘valleys’ define the phenotypic states that the cells can access. Rather than this deterministic picture, recent studies have discovered that cell fate is flexible and reversible, such that differentiated cells could be rejuvenated to a pluripotent state or directly jump from one ‘valley’ to the other^4,5^. As such, cell fate specifications can be interpreted as the mechanism by which cells make choices in the bifurcation landscape governed by the complex gene regulatory networks^6,7^. These networks determine the phenotypes of cells by controlling changes in gene expression through the activities of the transcription factors^8,9^, and extrinsic constraints like elaborate signalling dynamics and environmental cues^10,11^. Therefore, understanding the process of cell fate determination requires describing the integrative impact of both activities of transcription factors and extrinsic parameters on the cell fate decision landscape, as well as the underlying mechanisms.

It’s noteworthy that gene expression of transcription factors, besides specific regulation, also exhibits a strong dependence on the physiology of cells (e.g., cell growth rate). Prominent examples are found in both microorganisms and multicellular systems, such as the lysis-lysogeny decision of bacteriophage lambda infecting by fluctuations of gene expression rate coupled with cell physiological state^12,13^, the proliferation-to-invasion transition of *Bacillus subtilis*in response to nutrient limitations^14,15^, and epithelial-mesenchymal transition (EMT) controlling by surrounding niches^16-18^. It’s still a lack of direct evidence on how such global regulation of cellular physiology would introduce adaptations in gene regulatory networks and distort the cell fate decision landscape.

In this work, using *Escherichia coli*as a model system, we implemented a mutual repressive synthetic circuit^19-21^, characterized as a fundamental gene network that shapes a binary decision-making landscape^6,9^, to examine how its cell fate decision landscape would respond to growth variations and the accompanying dynamics of cell-fate determination. Pioneering bacterial physiology studies have demonstrated the global interdependence between gene expression and cell growth due to the re-allocation of cellular resources (e.g., building blocks, machinery for transcription and translation) and the adaption of metabolic networks^22-28^. Our results reveal that global coordination is not equal over the regulatory network. The unbalanced response in the two mutually repressive genes induces the bifurcating deformation of the fate-decision landscape. This reshaping process does not depend on specific molecular signalling or regulation but is an intrinsic characteristic of cell physiological regulation upon the fate determination networks.

## Results

### Growth-rate dependent phenotypic bistability

We introduced a well-established genetic circuit with a mutually repressive topology composed of two transcription factors (TFs), LacI (controlled by *P*_*L*_*tetO-1*promoter) and TetR (controlled by *Ptrc2*promoter)^19-21^into *E. coli*cells. This system can display two mutually repressive states by two distinct reporters, the green fluorescent protein GFPmut2 (GFP) and the red fluorescent protein mCherry (RFP), respectively (**Extended Data Fig. 1a**). During steady-state growth in a nutrient-rich growth medium (e.g., SOB, see **Methods**), cells can be induced to two distinct phenotypes (i.e., green state: high LacI/GFP expression; red state: high TetR/RFP expression) by the respective inducer, and maintain the same state for a long time even after removal of inducer (**Extended Data Fig. 1b-c**). This bistable phenomenon is known as the hysteresis of the mutually repressive system.

Surprisingly, we noticed that the phenotypic state cannot be always maintained during the batch culture growth. For a population initialized in the green state, both GFP and RFP intensity increased at the later log phase when the growth rate slows down. Further decrease in growth rate promoted a developmental progression where the GFP signal dropped abruptly with a concomitant increase of the RFP signal. We followed this state transition dynamics by defining the phenotypic changes during nutrient depletion (**Extended Data Fig. 2**). The ratio of cells in the green state monotonically decreased throughout a cell culture, as the growth shifted from a balanced-growth phase to a nearly stationary phase (**Fig. 1a**). Eventually, all cells initially in the green state shifted to the red state, when the growth rate dropped accompanied by nutrient depletion (**Fig. 1a**upper panel). Time series of single-cell fluorescent images over the growth curve more explicitly confirmed the phenotype transition from green state to red state (**Fig. 1b**). On the contrary, cells with an initial red state do not exhibit the state transition (**Fig. 1a**bottom panel). We confirmed that this transition is not a genotypic variation, as re-incubation of the cells from the stationary phase to the fresh SOB medium can recapture the bistability (**Extended Data Fig. 2**). Furthermore, cells incubated in the fresh SOB medium supplemented with conditioned medium collected from the stationary phase did not change their states, suggesting that the metabolic products generated during nutrient depletion do not contribute to the transition of cell phenotypes. Based on these observations, we hypothesized that the global physiology variations may underlie the reshaping of the cell fate decision landscape, implicating in the phenotype transition during growth rate decreases in batch culture growth.

**Fig. 1:**
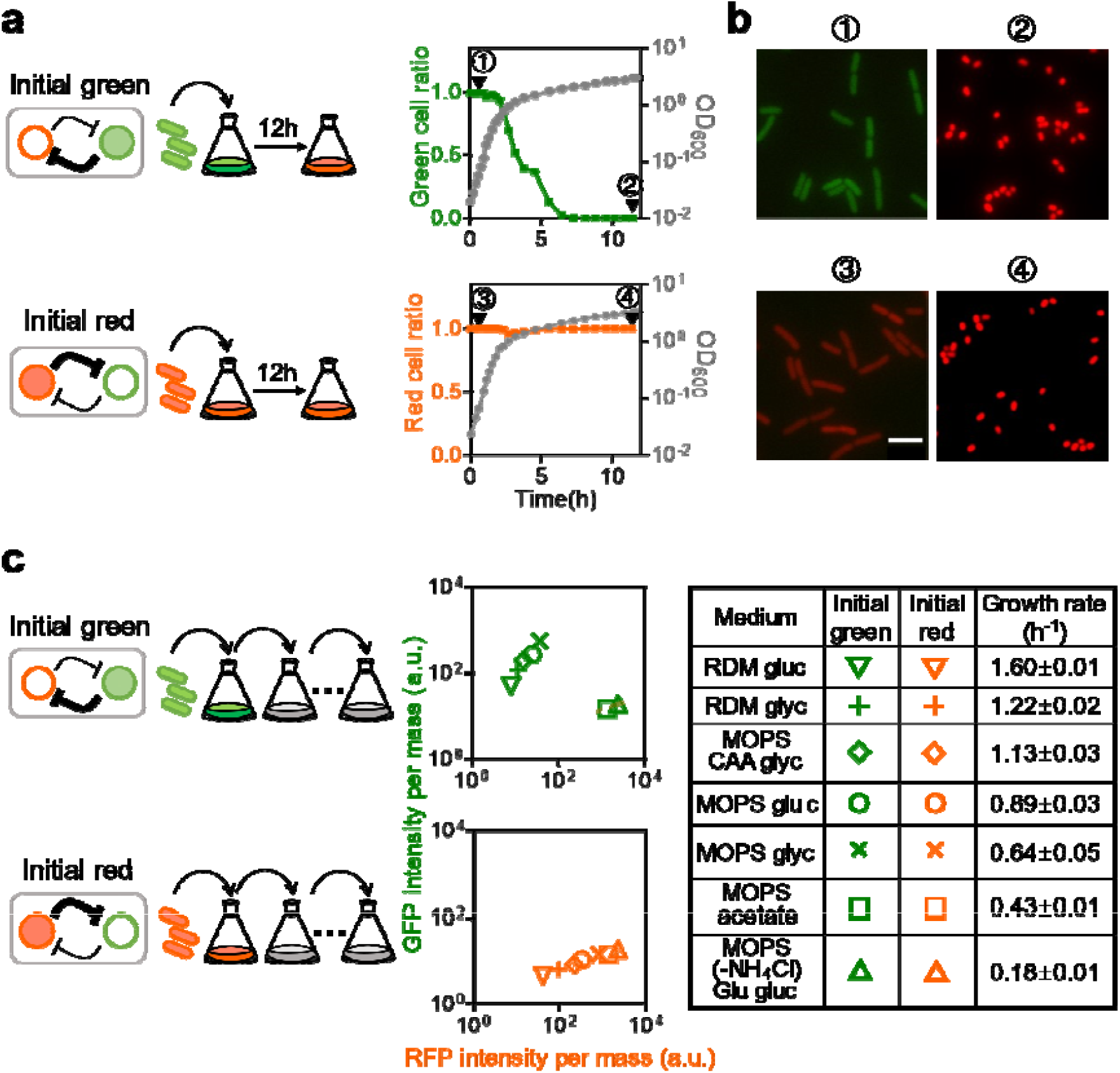
Cellular physiology-dependent state determination of a mutually repressive synthetic circuit. **(a)**The cell state kinetics during bulk culture growth, with cells initialized as two mutually repressive states: high LacI/GFP and low TetR/RFP (green state), or low LacI/GFP and high TetR/RFP (red state). To better discriminate the phenotypic states, we normalized the fluorescent signals by a separatrix in the scatter plot of GFP intensity per mass against RFP intensity per mass generated at each time point throughout the cell culture and thereby defined the cell ratio in the green state (**Extended Data Fig. 2**and **Supplementary Table 5**). Starting with cells in the green state (⍰), the green cell ratio decreases (the red cell ratio increases, **Extended Data Fig. 2**) until all cells transit into the red state (⍰), as the growth rate drops due to nutrient depletion of bulk culture. Meanwhile, starting with cells initialized in the red state (⍰), the red cell ratio keeps constant during the bulk culture growth, so that cells maintain in the red state (⍰). **(b)**Representative microscopic images of cells at different times (⍰-⍰), the scale bar shows 5 µm in distance. **(c)**Cell phenotypic state under different steady-state growth conditions (the growth media are listed, see **Supplementary Table 3**and **4**), with two different initial states: initial green state (green symbols) or initial red state (red symbols). Steady-state protein concentrations are given by the fluorescence intensity per mass measured under exponential growth conditions (see **Methods**). Given the two distinct states, we washed the cells and transferred them into a fresh medium without an inducer. Successive dilutions were carried out until the fluorescence intensity level varied less than 5% for each cell population so that the cells were maintained in steady-state growth. The mutual repressive circuit loses its bistability in slow-growth conditions (*λ*< 0.5 h^−1^).

Therefore, to validate the dependence of phenotype variations on cellular physiology, we examined the cell stable states under balanced exponential growth in different defined media to achieve various growth rates over an extensive range (**Supplementary Table 3**). In these conditions, the global effects of physiology are reflected first and foremost by the growth rate^22,24,25^. Cells were first fully induced to two respective states and kept growing exponentially via dilution regularly into fresh medium until steady state (**Fig. 1c**, details in **Methods**). Cells initialized as a red state can maintain the high red expression state over a wide range of growth rates (Growth rate *λ*from 1.60 h^-1^to 0.18 h^-1^, **Supplementary Table 6**), with the fluorescent protein concentration increases as the growth rate decreases (red symbols in **Fig. 1c, Supplementary Table 6**). However, cells starting with high GFP expression can only maintain the state from fast-to moderate-growth conditions (*λ*from 1.61 h^-1^to 0.59 h^-1^, green symbols in **Fig. 1c, Supplementary Table 6**). When steady-state growth rate was inferior to approximately 0.5 h^-1^, all green state cells switched to red state (green square and green triangle in **Fig. 1c**), suggesting the loss of bistability. We further demonstrated that the system exhibits a large range of hysteresis in a fast-growth medium (**Extended Data Fig. 1e**), but almost null of hysteresis in slow-growth conditions (**Extended Data Fig. 1f**), thereby confirming the loss of bistability. Additionally, we observed similar growth-rate dependence by growth rate titration via the translation-inhibiting antibiotic (**Extended Data Fig. 3**). The interplay between the growth-mediated global effects and specific mechanisms of the regulatory network may invoke unintended interactions, e.g., the growth modulation of gene expression^23^, growth feedback^29^, and growth retardation caused by metabolic burden^30^. Our experiments showed no significant difference in growth rates between the two distinct phenotypes (**Fig. 1c**right panel, **Supplementary Table 6**). Thus, the variation of phenotypes for this mutual repressive system is not induced by the nonlinear dilution due to the metabolic burden^30^caused by gene expression. These results lead us to expect that, the cell fate determination might take place without complex regulation and biochemical signalling.

### The unbalanced response of gene expression to growth-rate variations

To further elucidate how cellular physiology is involved in reshaping the fate decision landscape, we quantitatively examined the growth-rate-dependent global effects on gene expression of the mutually repressive circuit. We separated the circuit into two constitutively expressed parts (i.e., *Ptrc2*-driven *tetR*and *P*_*L*_*tetO-1*-driven *lacI*, **Extended Data Fig. 4a**) and characterized the corresponding expression capacities^31^(denoted as 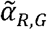in **Extended DataFig. 5**, defined as the constitutively expressed protein concentration under balanced exponential growth) by quantifying the fused fluorescent signals under various growth conditions. As shown in **Fig. 2a**, the expression capacities exhibit a strong growth rate-dependent effect. Both expression capacities increase faster than 1/*λ*as the growth rate decreases (**Extended Data Fig. 4b**), indicating that additional global regulation other than the cell volume variations in changing growth rate could be significant. Notably, the ratio of expression capacity at different growth rates, 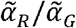, indicates the unbalanced variation of two proteins in response to the growth rate changes (**Extended Data Fig. 4c**): an approximately doubled increase in the ratio from fast-to slow-growth conditions. We further validated the unbalanced variations of two repressive genes by examining the mRNA abundance. The transcriptomics (mRNA-sequencing, see **Methods**) confirmed unequal dependence on growth rate in the expression capacities of two regulators (**Extended Data Fig. 4d-f**). Using an empirical dependence of translation activity on growth-rate^22-24,26^, we calculated the effective protein synthesis rates (denoted as α_*>R,G*_in **Extended Data Fig. 5**). The theoretical prediction can recapture the experimentally measured protein synthesis rates, of which there are maximums around the moderate growth rates (*λ*_*m*_≈ 0.5 h^-1^for *α*_*R*_and *λ*_*m*_≈ 0.75 h^-1^for *α*_*G*_; **Extended Data Fig. 4f-g**). Therefore, although the expression capacities and synthesis rates of both sides in the mutually repressive circuit show the sametrend of dependence on the growth rate, their detailed degree of dependence are not equal, which would break the balance of mutual repression.

**Fig. 2:**
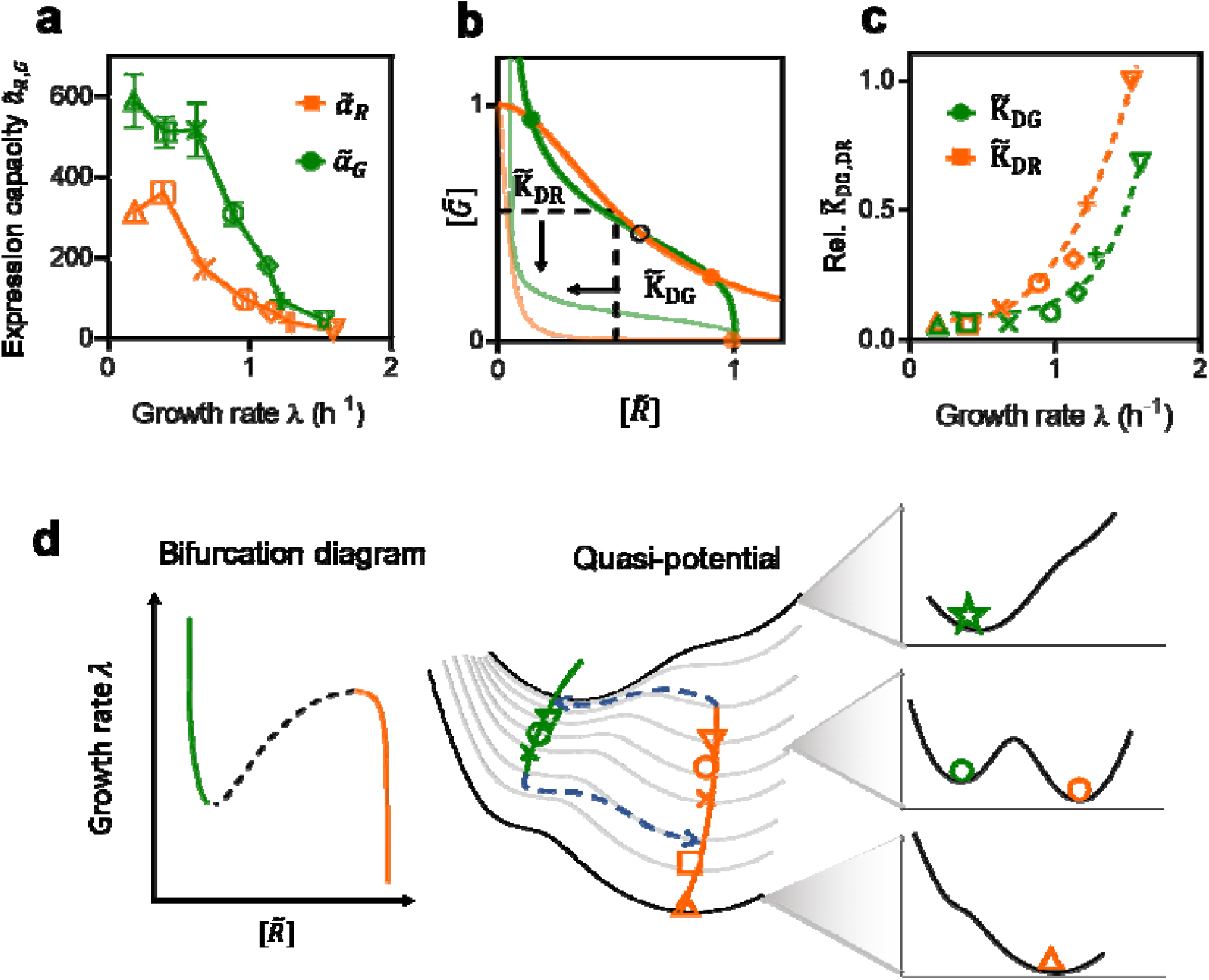
Cell growth-induced bifurcation and cell fate decision. **(a)**Growth-dependent expression capacity of constitutively expressed TetR (red symbols) and LacI (green symbols), respectively. Two separated parts that constitutively expressed *tetR*and *lacI*with C-terminal *sfgfp*fusion were used for the measurement of their corresponding expression capacity (**Extended Data Fig. 4a**). Different symbols correspond to growth media listed in **Fig. 1c**. Data represent the mean±SD for at least three biological replicates (**Supplementary Table 7**). **(b)**Illustration of the fixed points and their stability by nullclines plot for the mutually repressive system. The unbalanced variations of the dimensionless repression thresholds and result in the two nullclines experiencing a transition from three fixed points (by two solid lines, with two stable states marked as solid dots and one unstable saddle point marked as an open circle) to a single point (by two transparent lines, with only one stable state marked as a transparent dot). **(c)**Relative repression thresholds (normalized to 0 to 1) in units of gene expression capacity as a monotonically increasing function of growth rate. The repression thresholds are given in **Supplementary Table 9**and the gene expression capacities are indicated in **Fig. 2a**, and **Supplementary Table 7**. Symbols are experimentally calculated data, and the dashed lines indicate the model-fitted values (**Extended Data Fig. 4g**and **Supplementary Note 2**). **(d)**Schematic of quasi-potential landscapes under different growth rates. Cell fate decision is coordinated through growth-rate-dependent parameters by invoking growth rate A as the control parameter in a bifurcation diagram (left panel, solid lines represent the ensemble of stable steady-states, the dashed line indicates the ensemble of unstable steady-states, i.e., the saddle points). The evolution of the calculated quasi-potential landscape (see **Supplementary Note 4**) in varying the cell growth rate is given in the middle panel (symbols represent different conditions indicated in **Fig. 1c**). Different one-dimensional quasi-potential landscapes taking various growth rates from 1.6 h^-1^to 0.2 h^-1^were provided in **Supplementary Fig. 2**. It should be noted that our experimental conditions can only access to growth rate from 1.60 h^-1^to 0.18 h^-1^where the system’s state changes from bistable to monostable (only R state), thus we illustrate here the putative conditions when the system displays G state monostability (A > 1.60 h^-1^). At the critical growth rate, a saddle-node bifurcation takes place (dashed blue lines in the right panel) when one stable steady state is destabilized and shifts to the other one.

To study the consequence of these unbalanced variations of gene expression under different growth-rate conditions, we constructed a model to describe the mutually repressive circuit, considering the growth-dependent global gene regulation (**Supplementary Note 1**, and **Extended Data Fig. 5**). The model predicts that the stable steady states of the system are determined by the intersection of two nullclines^32^(solid lines in **Fig. 2b**) defined by the repressive regulatory functions. By nondimensionalized analysis, we found the dimensionless repression thresholds (**Supplementary Note 1**, normalized by their expression capacities under different growth rates and denoted as 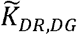) of the nullclines decrease steadily as growth rate decreases (**Fig. 2c**). However, the growth-rate-induced shifts are not balanced for the two dimensionless repressive thresholds. As a result, the number of possible intersections of the two nullclines changes in different growth rates, yielding the variations in available states of the system (from bistable to monostable, from solid lines to transparent lines as illustrated in **Fig. 2b**). Our analysis suggested that although it has been demonstrated that this mutually repressive circuit is robust to growth-mediated dilution^29^, the unbalanced dependence of gene expression on growth rate would impose a phase transition of available phenotypic states on the system (**Extended Data Fig. 6**)^23^.

The phase transition can be more explicitly illustrated by the quasi-potential landscape (**Fig. 2d**, and **Supplementary Note 4**), of which the local minima represent the stable phenotypic states of the system^33,34^. The calculated quasi-potential landscape exhibits a single minimum with highly expressed TetR/RFP in slow-growth conditions. As the growth rate increases, the landscape bifurcates into two local minima (the original one in the red state and the other new minimum in the green state), suggesting the system becomes bistable and hysteretic. Further increase in growth rate would impose the other saddle-node bifurcation, effectively leading to destabilization of the red state. Therefore, the growth rate plays a role as a global control parameter^35^to induce two bifurcation points from a monostable state to a bistable one, and further to the other monostable state. The above experiments only captured one bifurcating point from the bistable phase of either a green or red state to the monostable phase of a single red state as the growth rate decreases (coloured symbols in **Fig. 2d**). Another bifurcation from a green monostable state to bistability is not biologically accessible in our system.

### The determination of the critical growth rate of bifurcation

Further analysis of the model suggests that the bifurcation points would shift as the variations in the repression thresholds (i.e., *K*_*DR*_and *K*_*DG*_), which is governed by the equilibrium dissociation constant of the repressor from its corresponding binding operator^36^. Here, we constructed several modified LacI binding sites to obtain different dissociation constants (**Fig. 3a**, and **Extended Data Fig. 7**). Such designs enable us to vary the repression threshold *K*_*DG*_without affecting the Hill coefficient (cooperativity) of the regulatory function (*H*_*G*_*(R*)). We quantified the repression thresholds *K*_*DG*_for the modified circuits (**Fig. 3a**). Compared with the original circuit LO1, the two modified circuits exhibit a smaller *lac*operator dissociation constant (i.e., higher LacI binding affinity to their binding sites).

**Fig. 3:**
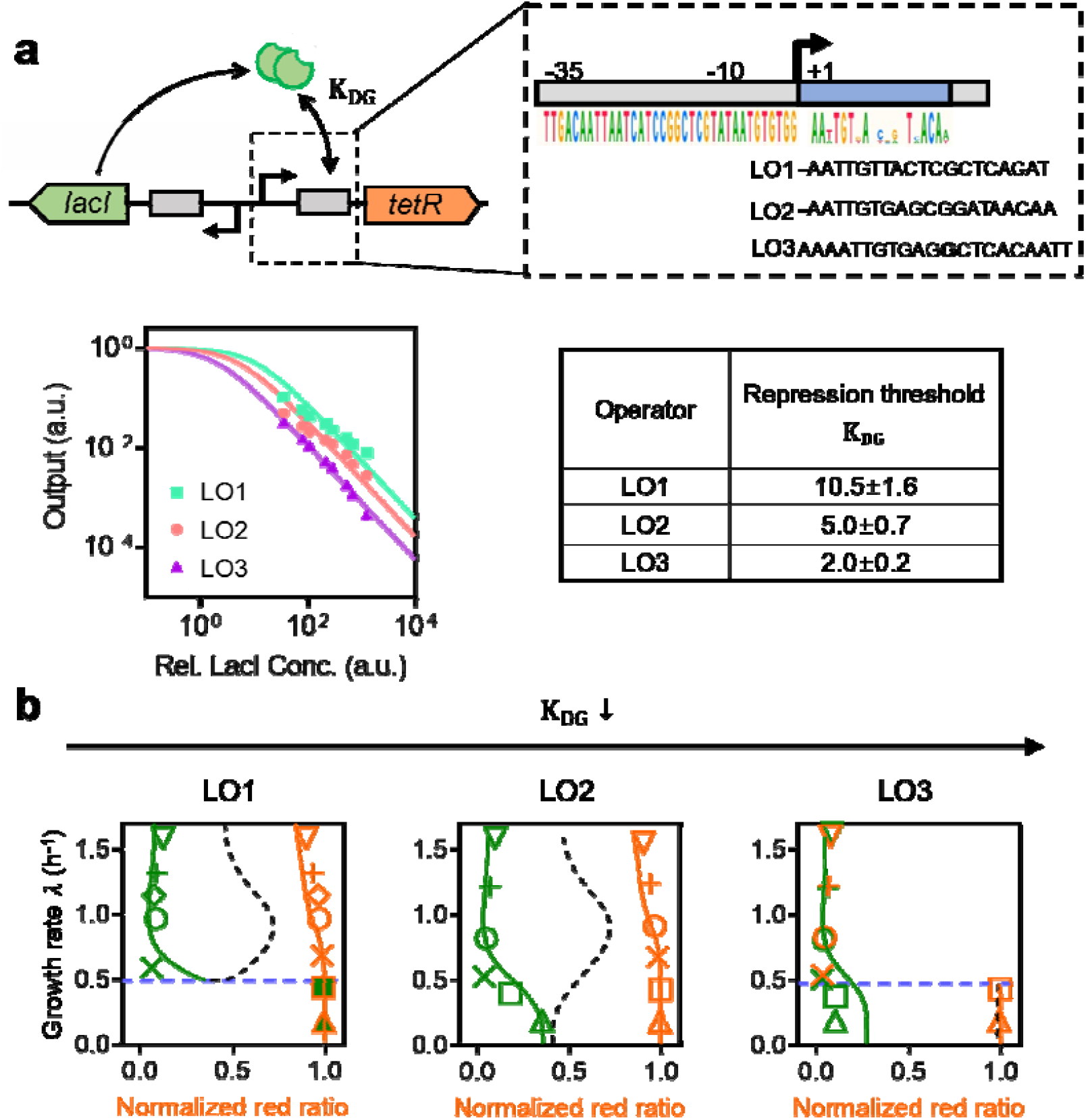
Titration of the repression threshold shifts the phenotypic bifurcation diagram. **(a)**Three representatives of synthetic binding sites for transcription factor LacI to vary the repression threshold . Quantification of repression response curves confirms the titration of the repression threshold without affecting the cooperativity of the repression response curves (**Extended Data Fig. 7**). The inputs and outputs are obtained by measuring the expression level of the fluorescence reporter (see **Methods**). Symbols represent the experimental data and solid lines donate parameter fitting lines with fixed Hill coefficient. The relative values of the repression threshold and standard deviation are provided in the table. **(b)**Bifurcation diagrams of mutually repressive systems with different repression thresholds, using growth rate as the key control parameter. The phenotypic state is quantified as the normalized red ratio, which is defined as the ratio of GFP fluorescent intensity to the sum of GFP and RFP fluorescent intensity under different growth conditions (**Supplementary Tables 6**and **8**). Different symbols are given in **Fig. 1c**. Solid lines indicate the ensembles of theoretically calculated protein concentrations at stable steady-state using the same normalization method as the experimental data, while black dashed lines point out the calculated saddle points that can hardly be measured experimentally. The quasi-potential landscape under fast growth (A 1.60 h^-1^) and slow growth (A 0.18 h^-1^) are indicated for each of the designs in **Supplementary Fig. 3**.

We further characterized the bifurcation diagrams of the modified circuits under different growth-rate conditions. Consistent with the model prediction, the circuits with different repression thresholds exhibit a shift of bifurcation points (**Fig. 3b**). Specifically, compared with the LO1 circuit that bifurcates from a monostable red state to bistable as the growth rate increases (**Fig. 3b**LO1), the circuit with a smaller dissociation constant *K*_*DG*_(i.e., with higher binding affinity) can maintain bistability over the wide range of growth rates (**Fig. 3b**LO2), suggesting it’s a robust design for a “toggle switch” regardless of variations in growth rates. Further reduction of *K*_*DG*_leads the system to become monostable (green state) for the fast growth rates (*λ*> 0.5 h^-1^) and bistability under *λ*< 0.5 h^-1^(**Fig. 3b**LO3). The abovementioned experiments as well as the model demonstrated that the growth-mediated gene expression could reshape the phenotypic potential landscape, yielding the transition of states through bifurcation points by varying growth rates.

### Growth-rate transition across the bifurcation point

Although the landscape deformation enables variations in phenotypes, it remains unclear how cell state is determined when the growth rate changes across the bifurcation points. According to Waddington’s metaphor of cell fate decision, the activity of regulatory networks shapes the landscape and determines phenotypic states, while fate decision trajectories are more likely to traverse during development under varying environments^3,35^. To address this question, we characterized the dynamics of cell phenotype determination across the bifurcation point during growth down- and up-shift, by monitoring the averaged fluorescent protein concentrations of initial green state populations (**Fig. 4a**, see **Methods**). Accompany by growth shifts, the cell state would change but along distinct trajectories depending on the landscape topology of the circuits (**Fig. 4b**).

**Fig. 4:**
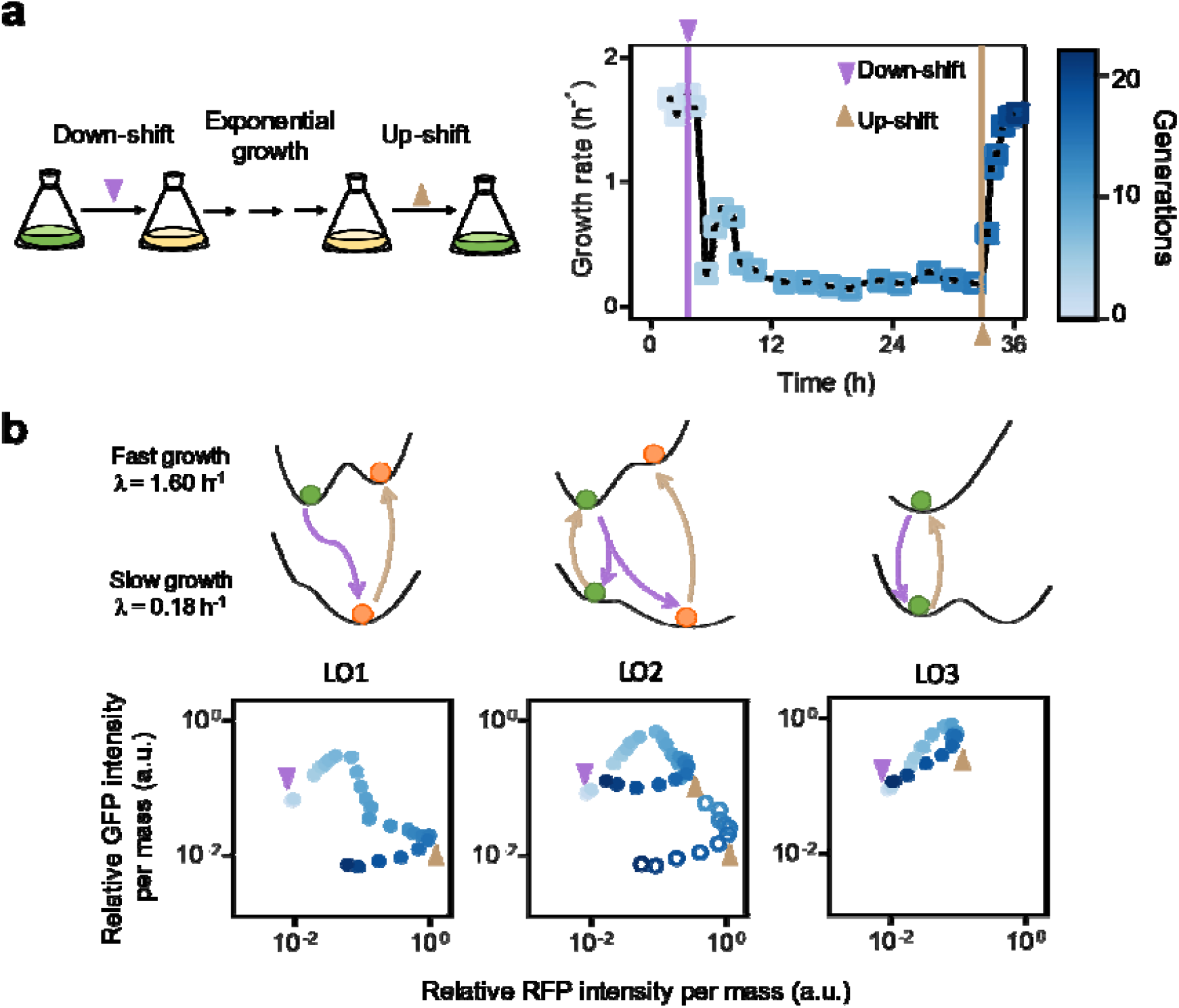
Irreversible and reversible cell fate determination. **(a)**Illustration of growth downshift from RDM glucose medium to MOPS (-NH_4_Cl) glutamate glucose medium and upshift experiments (see **Methods**), and the corresponding instantaneous growth rate. The violet down-arrow indicates the time at which the rich medium is depleted, and the yellow up-arrow refers to the moment the nutrient upshift is performed. The colours of symbols in the right panel correspond to the number of generations since t=0, which is determined by counting the doubling of cell mass (see **Methods**). After nutrient downshifts, the instantaneous growth rate *λ*drops abruptly, followed by rapid recovery, and overshot until reaches around 0.9 h^-1^and then further slows down towards the steady-state growth rate (∼0.18 h^-1^) in the after-shift medium. During the growth upshift, the instantaneous growth converged to the final rate in several hours. The cell growth dynamics are not affected by the different genetic circuits mentioned above, for the kinetics of cell growth were highly consistent among different strains, see **Extended Data Fig. 8. (b)**Trajectories of cell fate determination during growth shift. The quasi-potential landscapes for fast (1.60 h^-1^) and slow growth (0.18 h^-1^) conditions are provided in the top panels (**Supplementary Fig. 3**, we zoom in on the calculated landscapes for clarity). Population-averaged protein concentrations are plotted in a LacI-GFP versus TetR-RFP plane with colours indicating the numbers of generations in the bottom panel. The solid circles and hollow ones refer to two subpopulations characterized. Cells exhibit irreversible fate determination through saddle-node bifurcation (LO1, and LO2 hollow circles) and reversible ones (LO2 solid circles, and LO3) as well.

For the LO1 strain which has bistability at fast growth yet only the red monostable state allowed at slow growth, cells initialized as the green state at fast growth condition experience a phenotypic state transition during growth downshift. For the first several generations after the downshift, both GFP and RFP concentrations increase (**Fig. 4b**LO1), because of a higher protein synthesis rate with a slower growth-mediated dilution of protein throughout the growth downshift. The further increase in TetR/RFP expression triggers the repression of LacI/GFP, promoting a temporal progression of the cell state where the GFP signal drops with a concomitant increase of the RFP signal until the steady state. The sequential growth upshift starting with cells in the red state maintains the high TetR/RFP expression, suggesting the hysteresis and irreversibility properties. Alternatively, the LO3 strain retains its initial green state throughout the down- and up-shift (**Fig. 4b**LO3). During the growth downshift, both GFP and RFP signals of LO3 strain increase and eventually reach the steady state where GFP was highly expressed. Meanwhile, during the growth upshift, the cell exhibits a reduction of both fluorescent signals, but not a complete reversible trajectory as that of the downshift. The reduction rate of both GFP and RFP signals is fast initially and decays, exhibiting a strong correlation with the kinetics of growth rate^27^. Similar correlated dynamics can also be observed in LO1 strain with an initial red phenotype during the growth down- and up-shift (**Extended Data Fig. 8**). With a quasi-equilibrium assumption that the effective synthesis rates 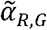are determined by the instantaneous growth rate *λ*(*t*), our mechanistic model can recapture the main dynamic features of the state transition driven by growth shifts (**Extended Data Fig. 9**, and **Supplementary Note 3**). Therefore, besides the direct constraint by the defined regulatory network, the dynamics of cell state transition also depend on the kinetics of growth transition.

Our deterministic model integrating the gene regulatory network with the global physiological constraints provides mechanistic insight into how growth variations could reshape the landscape and navigate cell-fate determination. However, this deterministic interpretation still encounters several challenges to describe the details of cell state transition dynamics during growth shifts. For example, the growth shift between two bistable landscapes without bifurcation (e.g., LO2 strain) would induce the bimodal gene expression (**Fig. 4b**LO2). During the growth downshift, the population of LO2 strains in the green state first went through a co-activation of both side genes of the mutual repression, and then split into two groups: one stays in the co-activation of both genes (dots in **Fig. 4b**LO2), and the other one in high TetR/RFP state with LacI/GFP repressed (circles in **Fig. 4b**LO2). We hypothesized that the co-activation state is because the potential landscape has a shallow barrier (**Fig. 4b**LO2), such that noise in the gene network could aid and trigger the hopping between the two “valleys”, resulting in a bimodal distribution of the gene expression level in an isogenous population^37,38^. Meanwhile, the following growth upshift drove the two groups of cells into mutually repressive states where only one side of the gene is highly expressed. Further detailed examination of single-cell lineage tracing^39-41^is required to address how and to what extent the stochasticity of the gene expression affects the cell commitment during the landscape deformation induced by growth shift^38^. Here, with experimental relevant parameters of noise, we introduced a stochastic scene into our model, thereby recapturing the potential landscapes^42,43^(**Supplementary Note 5**, and **Extended Data Fig. 10**). The barrier height in the landscape defines the transition feasibility between the two states^42^(**Extended Data Fig. 10d**, see **Methods**). Comparing the relative changes in barrier heights, the ratios of the barrier heights^44^are higher at slow growth rates (**Extended Data Fig. 10d**black solid lines), indicating that the cells tend to stay at the R state at slow growth rates. This further explains the strain LO2 could differentiate into two populations of cells during the growth downshift starting with a population at the G state (**Fig. 4b**LO2).

## Discussion

Classical developmental studies consider the deformation of Waddington’s landscape that varies the number of cell fate choices is usually induced by the upstream signalling, thereby driving the dynamics of cell differentiations^45,46^. Cell differentiation always exhibits a correlation between cell-phenotype transition and growth-rate variations. Prominent examples include the transition between epithelial and mesenchymal cells (EMT)^16-18^and the proliferation-to-invasion transition of *B. subtilis*^47^. However, the cause-and-effect relationship between two processes is always hard to distinguish in these natural systems^48^. The reconstitution by synthetic biology provides a complementary but important approach to recapitulate the fate decision processes with a fundamental set of regulations and enables the quantitative examinations of key parameters under well-controlled conditions^49-51^. This study, using a simple mutually repressive circuit as an example, demonstrated that global regulation of growth rate on the gene expression would break the balance of the regulatory network, thereby distorting the phenotypic landscape and further shaping the cell-fate determination. It is noteworthy to revisit the role of growth variations in spatiotemporal cell differentiation^37,49,52^.

## Methods

### Strains and construction

All experiments were done using strains derived from *E. coli*K-12 NCM3722 background. The strain NCM3722 was a gift from Dr. Chenli Liu. Strains derived from the wild-type strain are listed in **Supplementary Table 1**. Modified strains are edited by Lambda-Red recombination (pSIM series plasmids^53^) with polymerase chain reaction (PCR) amplified homologous fragments marked with antibiotic resistance genes, and the drug marker was eliminated by Cre-mediated site-specific recombination. All mutual repression circuits were derived from pECJ3 plasmid (gift from Dr. James Collins^20,21^, Addgene plasmid # 75465). Genetic circuits are harbouring in a plasmid with ColE1 origin. Genetic parts, including promoter, ribosome binding sites (RBS) and coding sequence were amplified from pECJ3 by PCR. Small parts which less than 200 bp were assembled via oligos (GENEWIZ Guangzhou Lab). Purified DNA fragments were assembled by One Step Cloning Kit (Vazyme Co., Ltd. Cat. C113) or Golden gate assembly^54^(NEB, Cat. M0202 and R3733). Sequence contents of genetic parts are collected in **Supplementary Table 11**.

### Growth media

Quantitative experiments were performed in MOPS buffered defined medium^55^supplemented with the appropriate antibiotic, i.e., 10 µg/mL kanamycin (Aladdin K103024), to maintain the plasmids. The MOPS buffer contains 40 mM MOPS (Sigma-Aldrich V900306), 4 mM Tricine (Sigma-Aldrich V900412, adjusted to pH 7.4 with KOH), 0.01 mM FeSO_4_, 0.276 mM K_2_SO_4_, 0.5 μM CaCl_2_, 0.525 mM MgCl_2_, 50 mM NaCl, 1.32 mM K_2_HPO_4_and micronutrient mixtures (3 nM (NH_4_)_6_Mo_7_O_24_, 0.4 μM H_3_BO_3_, 30 nM CoCl_2_, 10 nM CuSO_4_, 80 nM MnCl_2_, 10 nM ZnCl_2_). Unless otherwise specified, our MOPS buffer contains 9.5 mM NH_4_Cl as the nitrogen source. Different growth rates were achieved by providing different compositions of nutrients including carbon and nitrogen sources. Detailed information on these media is provided in **Supplementary Tables 3**and **4**. Various concentrations of mupirocin (Sangon Biotech A606674) were added to attain different growth rates through translation inhibition. The chemical inducers, IPTG (Sigma-Aldrich I6758, isopropyl β-D-1-thiogalactopyranoside) and cTc (Aladdin C103023, Chlorotetracycline hydrochloride) were diluted to an appropriate working solution when necessary.

The nutrient-rich bacterial growth medium SOB (Sigma-Aldrich H8032, Hanahan′s Broth) contains 0.186 g/L KCl, 2.4 g/L MgSO_4_, 0.5 g/L NaCl, 20 g/L tryptone and 5 g/L yeast extract. Cloning and genetic modifications used LB medium (HuanKai Microbial 028324, LB Broth without sugar), which contains 5 g/L NaCl, 10 g/L Tryptone, and 5 g/L yeast extract.

### Cell culture and growth rate measurement

Unless otherwise specified, cell culture and growth rate measurements were carried out in 3 steps: seed culture, pre-culture and experimental culture. Strains were streaked in LB agar plates from glycerol stocks. Cells were incubated at 37 □ for 10-12 hours. 3-5 colonies were picked from the agar plate and inoculated into a 14 mL tube containing 3 mL LB medium supplemented with an appropriate antibiotic. The culture was performed in a shaker (220 r.p.m., 37 □, Shanghai Zhichu Instrument) for at least 3 hours (or longer for cells with a slower growth rate) as seed culture. At the pre-culture step, cells were transferred to a 0.22 µm filter and washed with a pre-warmed experimental medium (at least 3-fold the volume of the seed culture) to remove the remnants of the LB medium. Washed cells were diluted into the experimental medium with a starting OD_600_(optical density at 600 nm, which was measured using a spectrophotometer Genesys 10s, Thermo Fisher Scientific) at approximately 0.01. Successive dilutions were achieved once OD_600_reached 0.2 and repeated for several rounds to keep a balanced growth condition. Samples for further quantitative measurements were collected to an OD_600_less than 0.2. Unless otherwise stated, pre-culture and experimental culture steps were performed in a water-bath shaker (150 r.p.m., 37 □, Shanghai Zhichu Instrument) in a 29 mm × 115 mm test tube with no more than 10 mL medium. For the growth rate measurement, cells were kept in pre-culture steps for at least 10 generations to establish steady-state growth. Experimental cultures were started by diluting the pre-culture to an OD_600_of approximately 0.02. Five to eight points were measured within a range of OD_600_from 0.03 to 0.2. The growth rate (*λ*) was calculated by fitting the data to an exponential growth curve.

### Flow cytometry analysis

The fluorescence intensity distribution of each sample was measured using a flow cytometer (CytoFLEX S, Beckman Coulter Life Sciences) with excitation/emission filters 488 nm/525 nm (FITC) for GFP (as well as sfGFP and mVenus) and 561 nm/610 nm (ECD) for mCherry. The gain for FSC, SSC, FITC and ECD channels were set to 500, 500, 250 and 1500, respectively. The trigger level was set manually on SSC-H according to different growth conditions. The flow rate was 60 µL/min and at least 50,000 events were recorded. The detected event number per second should be kept below 3000 events/second.

### Characterization of fluorescence intensity per mass

Fluorescence intensity per mass was employed to monitor the relative protein concentration. 10 µL samples of each culture were diluted into 500 µL pre-cooled MOPS buffer containing 5 mg/mL kanamycin to arrest protein translation and cell growth, immediately. The GFP intensity per mass and RFP intensity per mass were calculated by customized parameters: 1000XFITC-H/FSC-H (GFP intensity per mass) and 1000XECD-H/FSC-H (RFP intensity per mass). FSC-H was used to normalize the effect of cell size change in different growth conditions as its good linear relation with cell volume and cell mass^56^. All samples were stored in ice for 18 hours before measuring for fully maturing the fluorescent proteins.

### Cell state determination and classification

The cell states in different growth media were classified by determining the fluorescence intensity per mass of GFP and RFP using flow cytometry under exponential growth. cTc or IPTG was added into the growth medium with the final concentration of 10 ng/mL or 0.2 mM to trigger the cells into a high LacI/GFP state or high TetR/RFP one. Cells were subsequently washed with a pre-warmed experimental medium to eliminate the inducers and diluted into fresh medium without inducers until steady state. In the case of LO1, the green state (LacI/GFP state) refers to the condition where the steady-state GFP intensity per mass is higher than that of RFP (from RDM gluc to MOPS glyc medium, in **Fig. 1c**and **Supplementary Table 6**). Conversely, the red state (TetR/RFP state) was determined when the steady-state RFP intensity per mass is higher than that of GRP (from RDM gluc to MOPS(-NH_4_Cl) Glu gluc medium, in **Fig. 1c**and **Supplementary Table 6**).

### Protein expression capacity and synthesis rate measurement

We defined the protein expression capacity as the constitutively expressed protein concentration under balanced exponential growth, which indicates the highest level when the protein expression is unregulated. The expression capacity of *Ptrc2*-driven *tetR*and *P*_*L*_*tetO-1*-driven *lacI*(**Extended Data Fig. 4a**) was determined using flow cytometry by measuring the fluorescence intensity per mass as mentioned above.

For constitutively expressed protein, synthesis rate α was evaluated as the product of the concentration of an unregulated protein *P*^*\**^(i.e., the expression capacity) and the balanced growth rate *λ*\*:

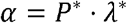

For instantaneous protein synthesis rate *α(t)*measurement, cells were collected and measured their relative protein concentration *(t)*and optical density OD*(t)*, simultaneously. The *α(t)*can be inferred by the equation:

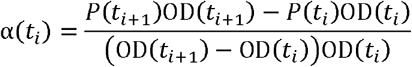

### Hysteresis experiment

Cells were kept growing exponentially for 10 generations (it takes usually ∼ 4.5 h in RDM glucose medium and ∼ 17 h in MOPS acetate medium) in the presence of 0.2 mM IPTG (to be triggered to red state) or 10 ng/mL cTc (to be triggered to green state). Cells were subsequently washed with a pre-warmed experimental medium to eliminate the inducers and diluted into a medium containing various concentrations of cTc within the range from 0 to 10 ng/mL for a further 10 generations. Samples were collected and detected using flow cytometer, 50,000 events were collected for each sample.

### Response curve measurements

All fluorescent intensity measurements were taken from flow cytometry. For characterizing the response function of the promoter variates, pRS3.R11_lacI series and pK4.5_C8 series plasmids were constructed. The pRS3.R11_lacI series plasmid harbouring the nahR-Psali expression system57 which can respond to the sodium salicylate (Sigma-Aldrich, S3007) was used to titrate the expression level of LacI, and the pRS3.R11_lacI-mVenus plasmid expressing lacI-mVenus fusion version was used to measure the LacI expression level in a specific concentration of the sodium salicylate, i.e., the input of the response curve (see **Extended Data Fig. 7**). Strain with blank plasmid pK4-Blank and pRS3.R11_mVenusn (mVenusn containing a C67Y mutation compared to wild type mVenus loses its chromophore) was used to measure the background fluorescent intensity. The promoter variates were inserted into the upstream of reporter gene mVenus in pK4.5_C8 plasmids for measuring the output of the response curve. The strains containing pRS3.R11_lacI and pK4.5_C8 plasmids were inoculated into RDM glucose medium supplied with proper antibiotics in 14 mL tube at 37 □, 220 r.p.m. for at least 8 hours. Overnight cultures were adjusted to OD_600_0.2 by diluting with MOPS buffer. After OD adjustment, cultures were diluted 150-fold into 149 μL RDM glycerol medium with antibiotics and specific sodium salicylate concentration. Cells were grown in a thermostatic microplate shaker (AOSHENG, MB100-4A) maintained at 37 □ and 1000 r.p.m., using flatten button 96-well-plates (Corning, 3336) and sealing with breathable film (Axygen, BF-400-S). After growing 7 generations, 15 μL cells were diluted into 135 μL pre-warmed fresh identical medium for 4 generations. 10 μL cultures were collected and diluted into 500 μL pre-chilled MOPS kanamycin buffer. All samples were stored in an ice slurry for 18 hours before the fluorescence intensity measuring.

### Nutrient-shift experiment

The downshift and upshift experiments presented in **Fig. 4**and **Extended Data Fig. 8**were carried out between RDM glucose medium and MOPS(-NH_4_Cl) glutamate glucose medium^58^. RDM glucose medium contains extra amino acids (EZ) and nucleotides (ACGU), which provides a fast growth rate. Before the growth downshift, the culture was in exponential growth at 37 □ in the RDM glucose medium. When OD_600_reached 0.15, a dilution was performed with a dilution ratio of 1/10 to the pre-warmed MOPS(-NH_4_Cl) glutamate glucose medium. Successive 1/5 dilutions to the same medium were applied once the OD_600_reached approximately 0.15 to keep the OD_600_of culture between 0.03 to 0.15. The OD_600_of the culture was measured frequently and an aliquot of cell suspension was taken at the same moment of OD_600_measurement for further flow cytometry analysis immediately. When cells’ fluorescence intensity levels varied less than 5% compared to the previous one, cells were considered to reach to steady state. A growth upshift was subsequently carried out to RDM glucose medium using the same procedure as described in growth downshift until steady state. The instantaneous growth rate was calculated as:

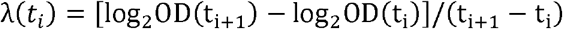

### Cell Imaging

Cells were imaged using Nikon Ti-E microscope equipped with 100X objective (NA=1.45). Illumination source was X-Cite 110LED, and Andor Zyla 4.2s sCMOS was used to capture images. Phase-contrast and widefield fluorescence images were acquired. Cell culture was adjusted to OD_600_0.1 before imaging. A 1.5% agarose pad with MOPS buffer was used to immobilize the cells. Inoculate 2 μL cell culture on a coverslip and cover by agarose pad gently. Each sample was collected and imaged within 5 min. For the green fluorescence channel, the emission filter and the excitation filter were FF02-482/18-25 and FF01-525/45-25 (Semrock), and the exposure time was 50 ms. For red fluorescence, emission and excitation filters are FF01-591/6-25 and FF01-647/57-25 (Semrock), and 100 ms exposure time.

### mRNA sequencing and data processing

Samples used for RNA extraction were prepared following the method described in the main text. About 1 OD/mL cells were mixed with 2 volumes of RNAprotect Bacteria Reagent (Qiagen, 76506), and the cell pellet was collected by centrifuging at 2500 g for 8 minutes at room temperature. For long-term storage, collected samples were stored at -70 °C. Cells were destructed in TE buffer containing lysozyme (Sigma-Aldrich, 62917) and incubated at room temperature for 5 min. After the lysis procedure, total RNA was extracted from cell lysate by RNeasy Plus Mini Kit (Qiagen, 74136). Total RNA which was carried out following the method described above was removed rRNA by rRNA Depletion Kit (Vazyme, N407-01), and cDNA library preparation and high-throughput sequencing were performed in MGI platform (BGI, Wuhan lab).

Sequencing raw data was processed using customed python scripts (https://github.com/MinTTT/RNA_seq_pip). Briefly, raw sequencing files were trimmed adapters using cutadapt^59^. Next, the sequencing data were aligned to reference sequences using Bowtie2^60^. The referencing sequences were generated according to the sample type, where the genome is NCM3722 (GenBank: CP011495.1) for all strains and the plasmids (pECJ3_lacI-sfgfp_delta_TetR or pECJ3_tetR-sfgfp_delta_LacI depending on the strains, the sequence details are listed in **Supplementary Table 2**). After alignment, reads coverage over the whole genome was calculated. Specifically, all properly mapped reads were separated according to their mapped strands using samtools61. Notice, we have removed the reads mapped to tRNA regions considering the bias caused by the non-uniform of tRNA in different samples^62^.

mRNA abundance 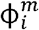was calculated by followed equation,

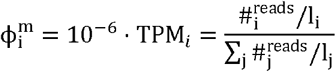

where 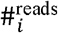represents the number of mapped reads of gene *i, l*_*i*_denotes the length of gene *i*.

### Mathematical modelling

Ordinary differential equation (ODE) models were developed to describe and analyze the growth-rate dependency of cell fate stability of the mutual repressive circuit at the population level (**Supplementary Note 1**). The growth-rate-dependent protein synthesis rate of different constitutive genes was formulated using the empirical relations and key assumptions of bacterial “growth law” (**Supplementary Notes 2**and **3**). The one-dimensional quasi-potential landscapes calculated from the dimensionless ODEs were to illustrate the variations in the system’s stability (**Supplementary Note 4**). The probability potential landscape was also calculated (**Supplementary Note 5**). The barrier heights as the potential difference between the local minimum and the corresponding saddle point, i.e., *U*_barrier =_*U*_saddle_– *U*_min_, for quantifying the robustness of each state.

## Data availability

Sequencing data have been deposited to the NCBI BioProject with the accession code PRJNA835246. Major experimental data supporting the findings of this study are available within the main text and Supplementary Information. Other data are available from the corresponding author on reasonable request. Custom-made simulation code is available via GitHub at https://github.com/Fulab-SIAT/cell_fate_2022.

## Acknowledgements

The authors thank C. Liu for sharing *E. coli*strain, X. Li, Y. Bai, S. Huang, C. Lou, F. Jin, C. Liu, C. He, Z. Ma for discussions, comments, and technical support.

## Author Contributions

Conceptualization: J.Z., P.C., and X.F.; Methodology: J.Z., P.C., and X.F.; Investigation: J.Z., P.C., and X.F.; Funding acquisition: J.Z., and X.F.; Writing – original draft: J.Z., P.C., and X.F.; Writing – review & editing: J.Z., P.C., and X.F.; Supervision: X.F.; Project administration: X.F.

## Competing interests

The authors declare no competing interests.

**Extended Data Fig. 1:**
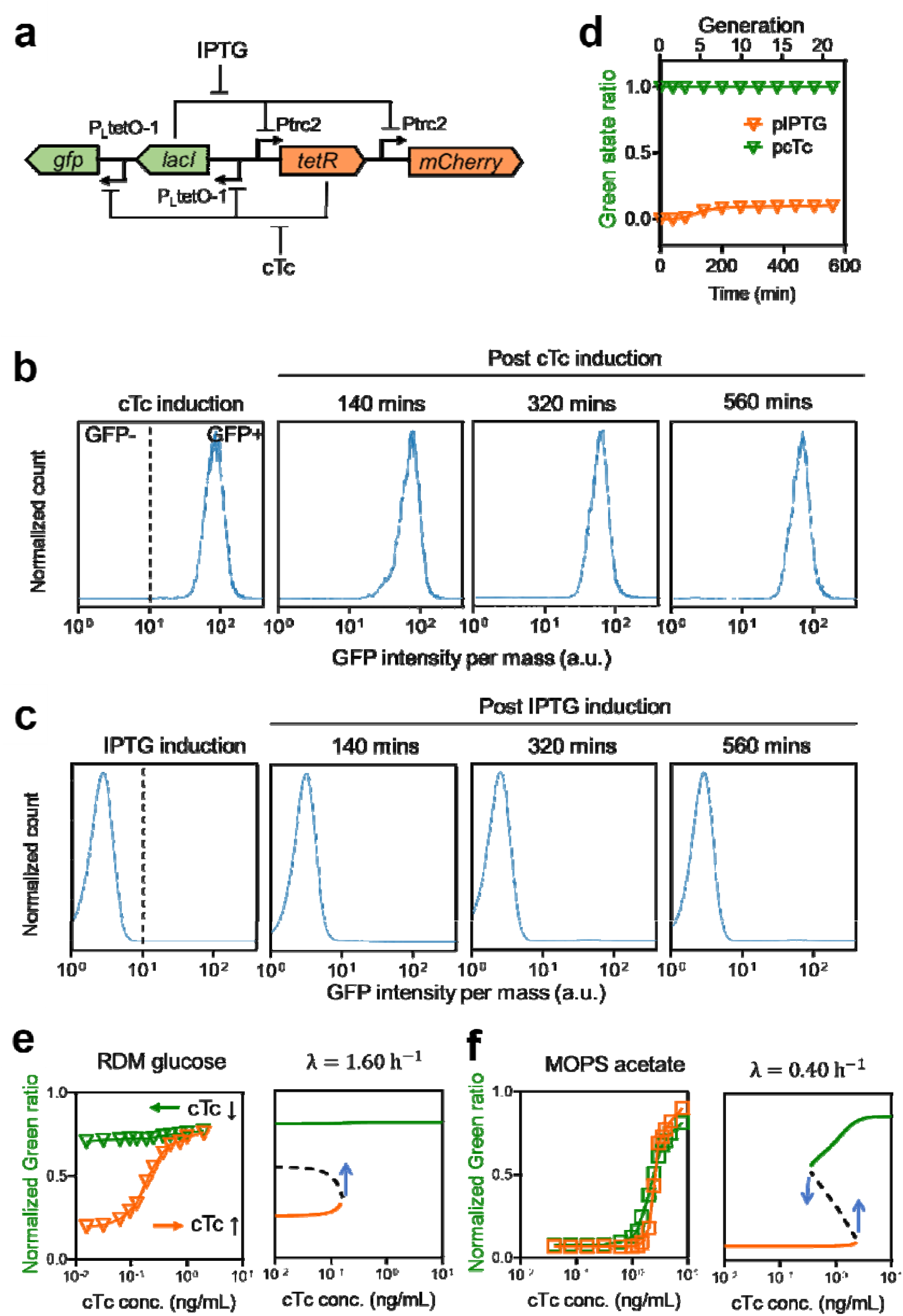
Schematic illustration of the design of the mutual repressive circuit and demonstration of bistability. **(a)**This circuit consists of reciprocal transcriptional repression by TetR and LacI. mCherry and GFP serve as reporters for TetR high state and LacI high state, respectively. The addition of cTc relieves TetR repression, allowing for high expression of LacI and GFP, whereas induction with IPTG relieves the LacI repression, allowing for high expression of TetR and mCherry. All mutual repression circuits discussed in this work were derived from pECJ3 plasmid (gift from Dr. James Collins, Addgene plasmid # 75465). **(b-c)**Temporal histograms of cell states under exponential growth in RDM glucose medium following prolonged culture via successive dilution to demonstrate the irreversibility of cellular states. Cells were induced with either 10 ng/mL cTc (**b**) or 0.2 mM IPTG (**c**) and kept growing exponentially for approximately 10 generations (∼ 4.5 h). Flow cytometry measurements were taken at different time points after cells were washed and transferred to a medium without the inducer (post-induction). Cells can be able to successfully maintain their corresponding phenotypes upon removal of the inducers. Owing to the fluctuations in gene expression, a small population (∼ 10%) of post-inducted red state cells (i.e., low GFP state) was observed to spontaneously switch to the green state (i.e., high GFP state) within 20 generations (as indicated in panel **d**). **(e-f)**Hysteresis experiments were carried out by measuring the dose-response curves in varying cTc concentrations under fast and slow growth conditions (see **Methods**). Red and green symbols represent initial red state cells and initial green state ones, respectively. **(e)**A large range of hysteresis was revealed in RDM glucose medium, suggesting a bistable behaviour under a fast growth rate. The bifurcation diagram calculated from the deterministic model in varying inducer concentrations with a given growth rate is shown. The red curves show the ensembles of red states, the green curves show the ensembles of green states, and the black dashed lines suggest the calculated saddle points which are unstable. The appearance of bimodal distributions in the range of 0.1 to 1.0 ng/mL cTc concentration is supposed to be the gene expression fluctuations in proximity to the bifurcation point (blue arrows in the right panel). **(f)**A loss of bistability was confirmed in MOPS acetate medium without inducer. The hysteresis experiment shows that the bistable range is significantly reduced in the MOPS acetate medium, within a small range of cTc level in the vicinity of 1 ng/mL, and this was confirmed by simulation.

**Extended Data Fig. 2:**
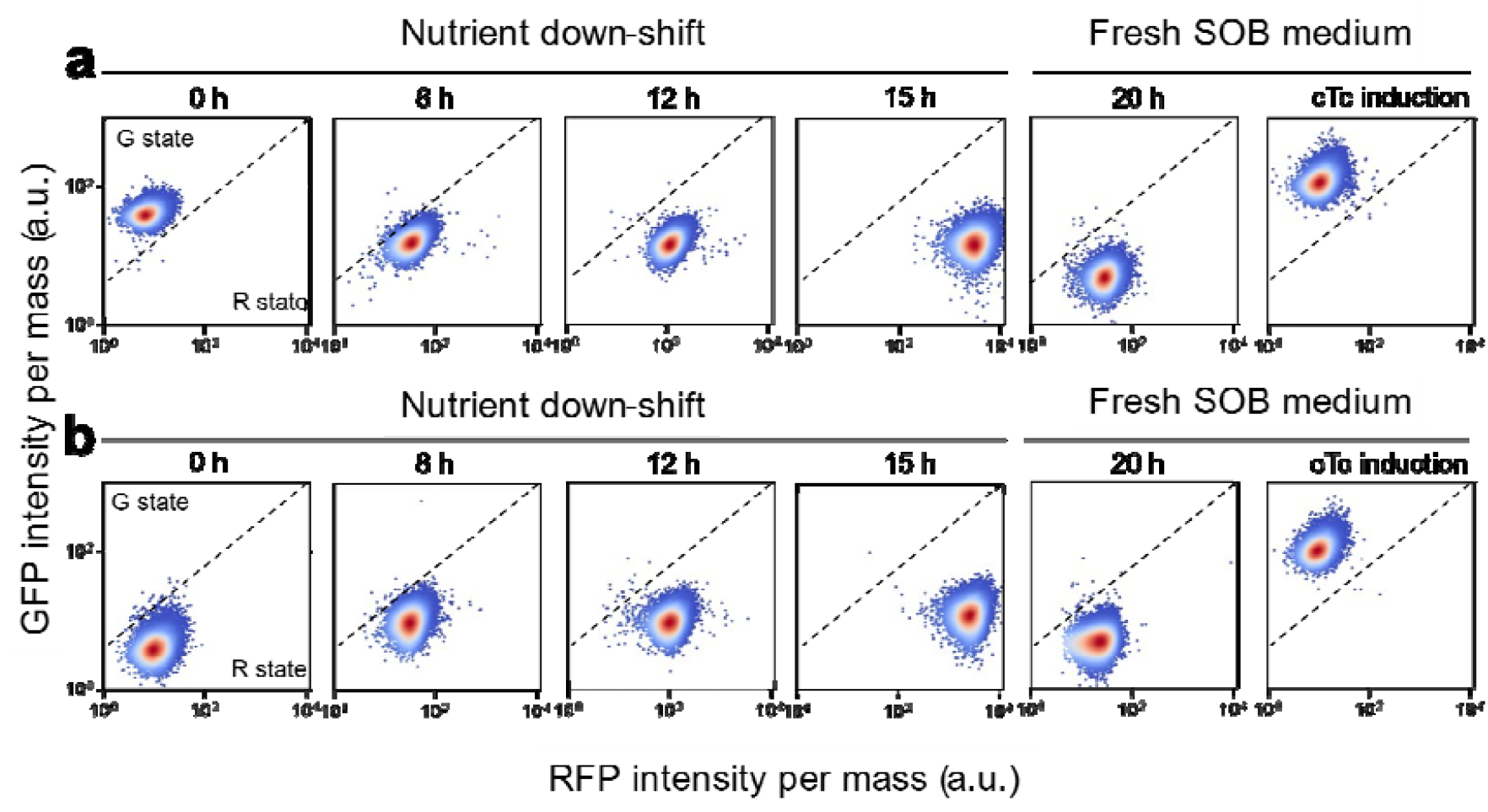
Temporal scatter density plots of cell states during nutrient depletion in SOB batch culture and re-incubation of the cells from stationary phase to fresh SOB medium. Data were obtained by flow cytometry at the indicated times with 50,000 events detected per experiment (**Supplementary Table 5**). A dashed line in the density plot is indicated to discriminate between two distinct phenotypic states (see **Methods**). (**a**) A time course is shown for cells initially induced to green state switching gradually from a green state (0 h) to a red state (15 h) throughout the nutrient depletion until the stationary phase. These cells maintained the red state when diluted into fresh SOB medium and grown on the rich-nutrient condition for another 5 h. (**b**) A time course for cells initially switched to the red state shows that they do not change their state during nutrient depletion. Cells can be further switched to the green state by adding cTc and maintaining this state in a no-inducer fresh medium. It shows that this circuit recaptures its bistability and suggests the phenotype changes during the nutrient depletion do not involve genetic variations.

**Extended Data Fig. 3:**
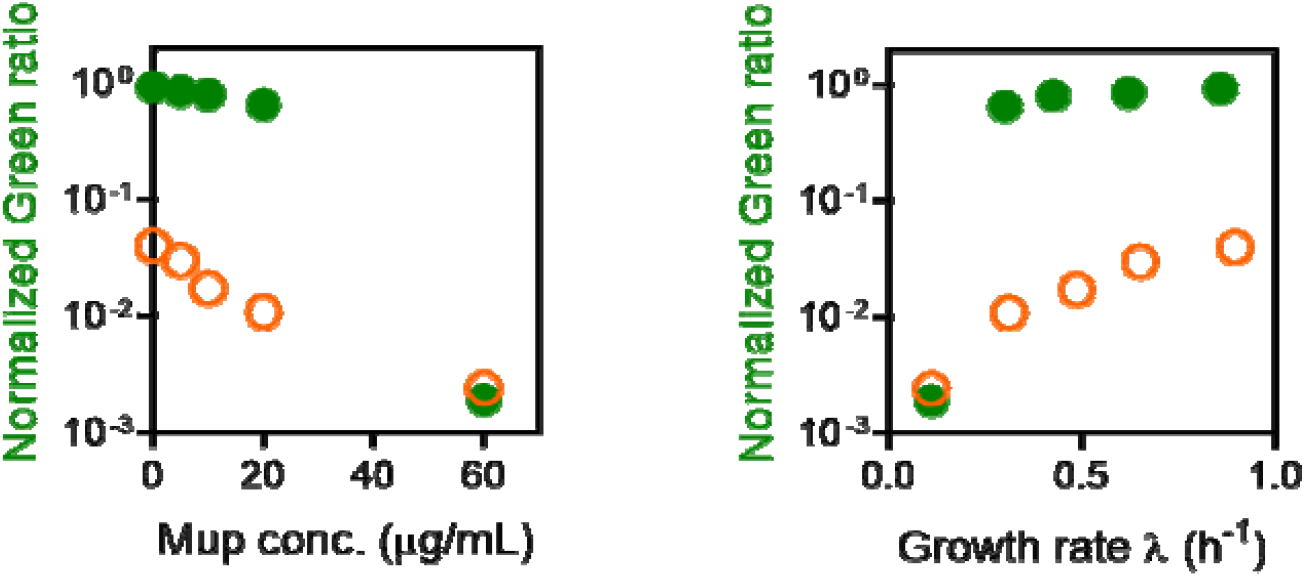
Variations of cell growth rate and stability of mutual repressive circuit under the treatment of diverse concentrations of sublethal dose of mupirocin. Mupirocin is a translation inhibitor that inhibits the charging of isoleucine tRNA, resulting in a reduction of the translational elongation rate of the ribosome. *E coli*. cells harbouring the mutual repressive circuit were grown in MOPS glucose minimal medium with diverse concentrations of mupirocin varying from 0 to 60 µg/mL. A loss of bistability was observed under slow growth when varying the cell growth rate by titrating the translational elongation rate.

**Extended Data Fig. 4:**
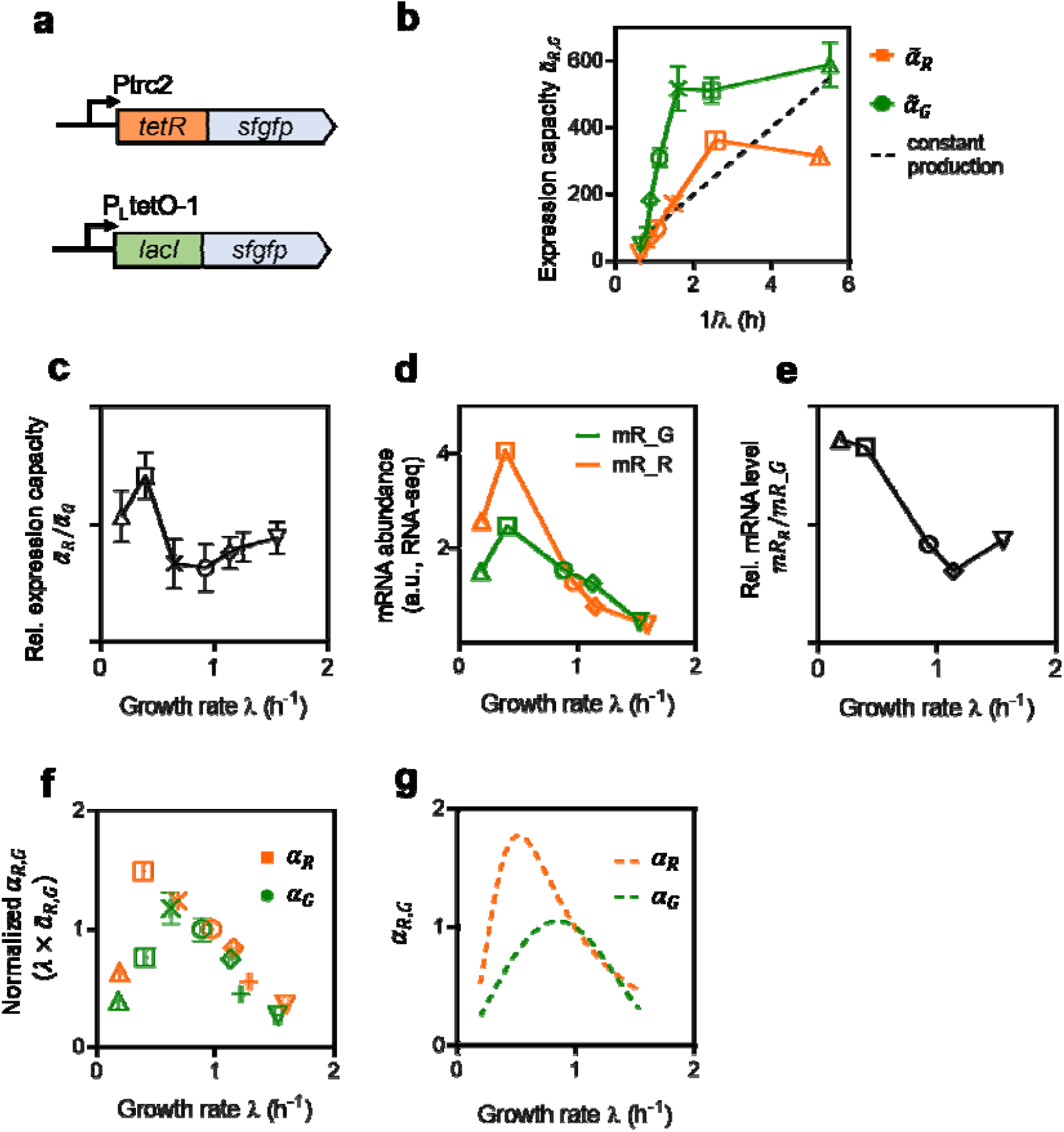
Growth rate dependence of constitutively expressed TetR and LacI. The protein synthesis rate is an effective value that includes the transcription rate, translation rate as well as the concentration of the corresponding gene and the degradation rate of the corresponding mRNA. For the regulatory system, protein synthesis is furthermore a function of transcription factors and other regulators related to cell physiology state. In this study, constitutively expressed *tetR*and *lacI*(panel **a**) were considered to separate the global effects of cell physiology state and specific regulations of TFs. (**a**) Two separated parts that constitutively expressed *tetR*and *lacI*with C-terminal *sfgfp*fusion were used for the measurement of their corresponding expression capacity. These two strains with the same reporter gene can eliminate the possible interaction between the mutually repressive parts, and thereby can represent the maximum expression levels that allow for each side of the mutually repressive circuit. (**b**) Expression capacities 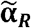and 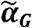of constitutively expressed TetR and LacI, respectively, as a function of **1**/**λ**. The dashed line indicates the steady-state expression capacity in the case of constant production of protein which is independent of growth rate (the constant value of protein production rate was the one of TetR in MOPS glucose medium). Both expression capacities increase faster than **1**/**λ**as the growth rate decreases. (**c**) Relative protein expression capacity is defined as the ratio of TetR and LacI protein concentrations under different growth conditions. Results derived from data shown in **Fig. 2a and Supplementary Table 7**. (**d**) Relation between the mRNA abundance and growth rate during steady-state growth. (**e**) Relative mRNA level, defined as the ratio of mRNA abundance of *tetR*and *lacI*genes under different growth conditions (results derived from data shown in **d**). (**f**) Growth rate dependence of the normalized protein synthesis rates ***α***_***R***_and ***α***_***G***_. ***α***_***R***_and ***α***_***G***_are derived from experimental data of tetR and lacI expression, respectively, under different growth conditions, i.e.,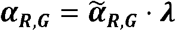. The synthesis rate obtained in different growth media was normalized to the value in MOPS glucose medium. Both synthesis rates have maxima around the moderate growth rates (∼ 0.5 h^-1^for ***α***_***R***_and ∼ 0.75 h^-1^for *α*_*G*_). The non-monotonic behaviour of ***α***along with growth rate could be explained by a trade-off between the availability of transcription and translation machinery(e.g., ribosome, RNAP etc.)^22,24^, translation elongation rate^26^, and modulation of plasmid copy number by replication control^63^. (**g**) Prediction of growth rate dependence of protein synthesis rate ***α***obtained from measured mRNA abundance in panel **d**by considering the empirical growth rate dependence of translation activity (see details in **Supplementary Note 3**).

**Extended Data Fig. 5:**
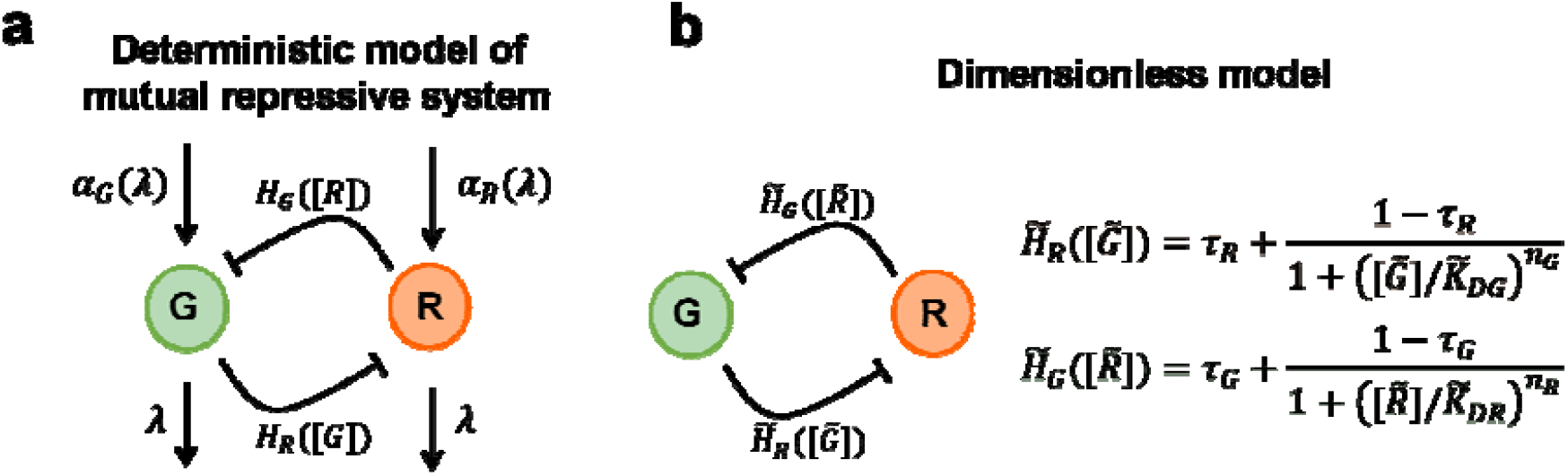
Dimensionless analysis of the deterministic model for the mutual repressive system. **(a)**The model includes the global regulation of cell growth along with growth-dependent protein synthesis ( and ) and degradation (growth dilution under the assumption that protein turnover is negligible) as input stimuli and output of the system (Supplementary Note 1). The model for the mutual repressive system is typically written as and, where and are the mutual repressive relations between TFs, which are described as two decreasing Hill functions of [R] (the reporter of TetR expression) and [G] (the reporter of LacI expression) with repression thresholds, respectively. **(b)**By defining the expression capacity, we would denote, as the dimensionless concentrations, and as the generational time of the system, thereby obtaining: — and —. Due to the growth rate dependence, the dimensionless repressive thresholds and would effectively increase with the growth rates but with different magnitudes. Therefore, the dimensionless nullclines of the system where and, such that and shifts as the growth rate changes, causing variations in the number of fixed points (**Fig. 2b**).

**Extended Data Fig. 6:**
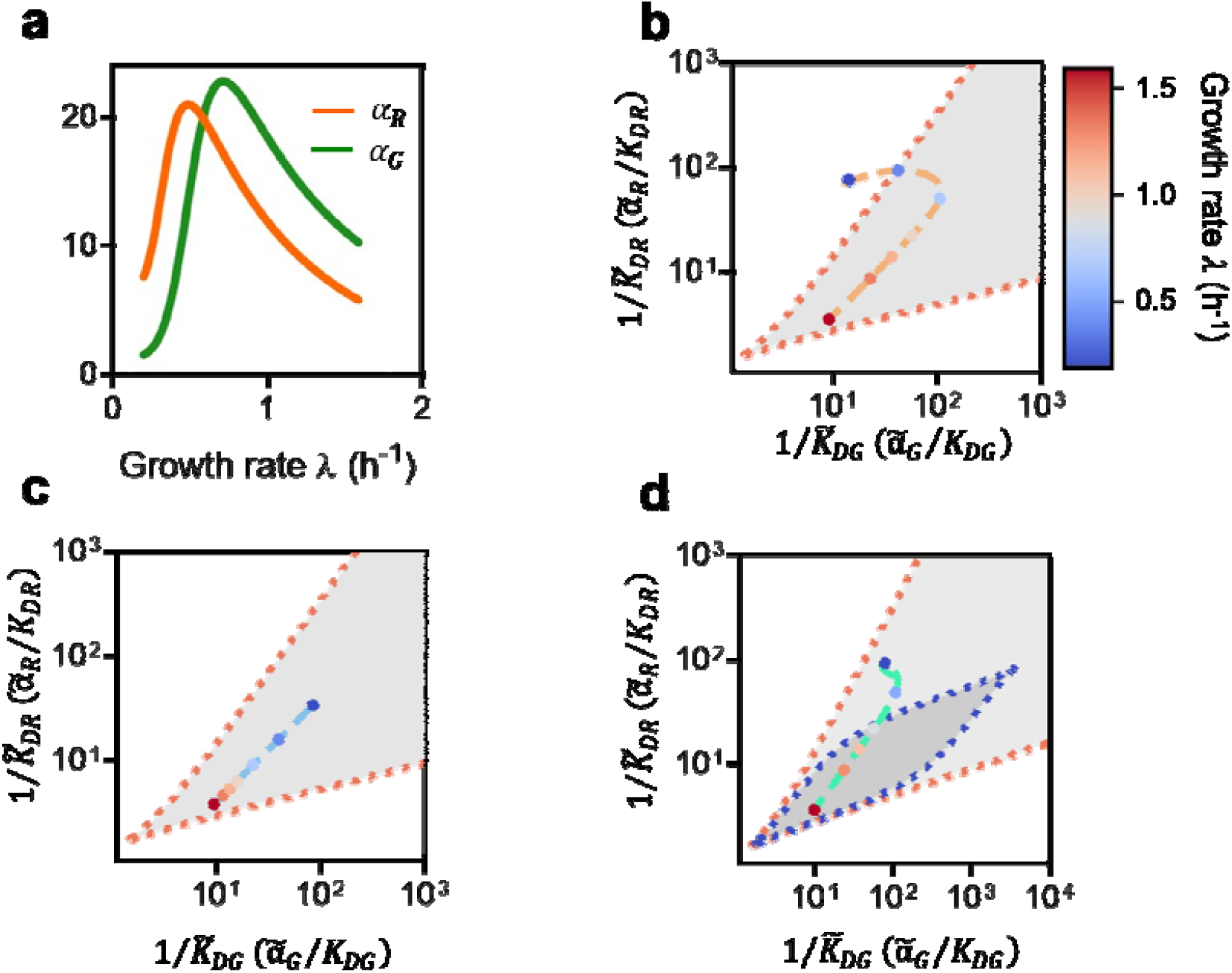
Growth-rate-dependent gene expression and promoter leakage enable phase transition in the bistable mutual repression circuit. We demonstrate that the growth-rate-dependent protein synthesis rate origins the bifurcation of the mutual repression circuit by mathematic modelling. (**a**) As the experimental data (**Extended Data Fig. 4f**) indicated, the protein synthesis rates,, are nonlinear to growth rate and shown in a context-dependent manner. (**b**) is the dimensionless repressive threshold which is linear to (**Supplementary Note 1**), and the bistability of the circuit is determined. We plot the phase diagram of the circuit with the promoters without expression leakage (i.e., both are equal to 0). When both stay at the grey region, the circuit exhibits bistability. Titrate the growth rate from 1.6 h^-1^to 0.2 h^-1^, both of the two mutual repression sides are varying and the trajectory of the circuit state indicates the bifurcation process (orange dashed line). Both protein synthesis rates are two orthogonal forces driving the system to traverse the phase diagram and lead to a long and curved trajectory than if is constant (**c**), resulting in bifurcation. (**c**) The trajectory of the strain (blue dashed line) with protein synthesis rates is constant and is plotted as a reference, which doesn’t enable the bifurcation. (**d**) The bistable region can be eroded by the leakage expression (blue dashed line encircled). We fix ***τ***_***R***_and ***τ***_***G***_are 0.035 and 0.002, respectively. The green dashed line is the trajectory that is fitted from experimental data (**Fig. 2a**).

**Extended Data Fig. 7:**
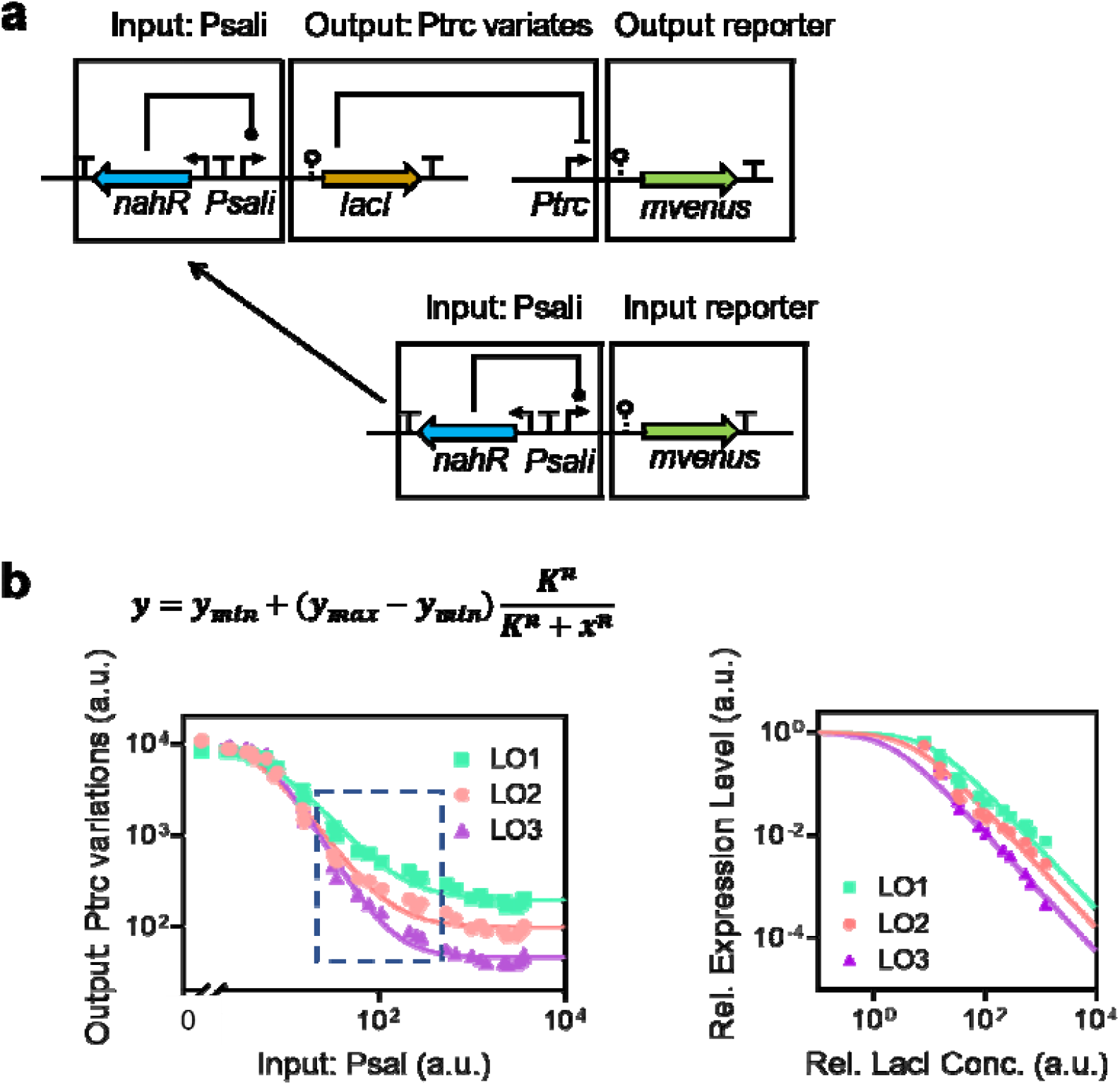
Measurement of response curves of the regulatory systems and parameter fitting. An experimental framework reported in previous work^64^was used for the characterization of *lac*repressor operators. **(a)**The expression level of this transcriptional repressor LacI is indicated via the intensity of fluorescent protein, mVenus, which is independently driven by the identical input promoter, i.e., *Psali*. The output of this regulatory system is indicated by the same reporter. By this means, the input-output response function could quantitatively describe the properties of the interaction between the *lac*repressor and its binding site. The plasmids used are given in **Supplementary Table 2. (b)**Experimental measurements, and parameter fitting of the response curves for different circuits. For each of the measurements, various concentrations of sodium salicylate were used to induce the *Psali*promoter activity. Input and output were both identified as fluorescent intensity per mass using flow cytometry (see **Methods**). Only a moderate range of data (within the dotted box in the left panel) was taken for parameter fitting. The Hill coefficient was fixed at 1.15 and other parameters were obtained using best-fit.

**Extended Data Fig. 8:**
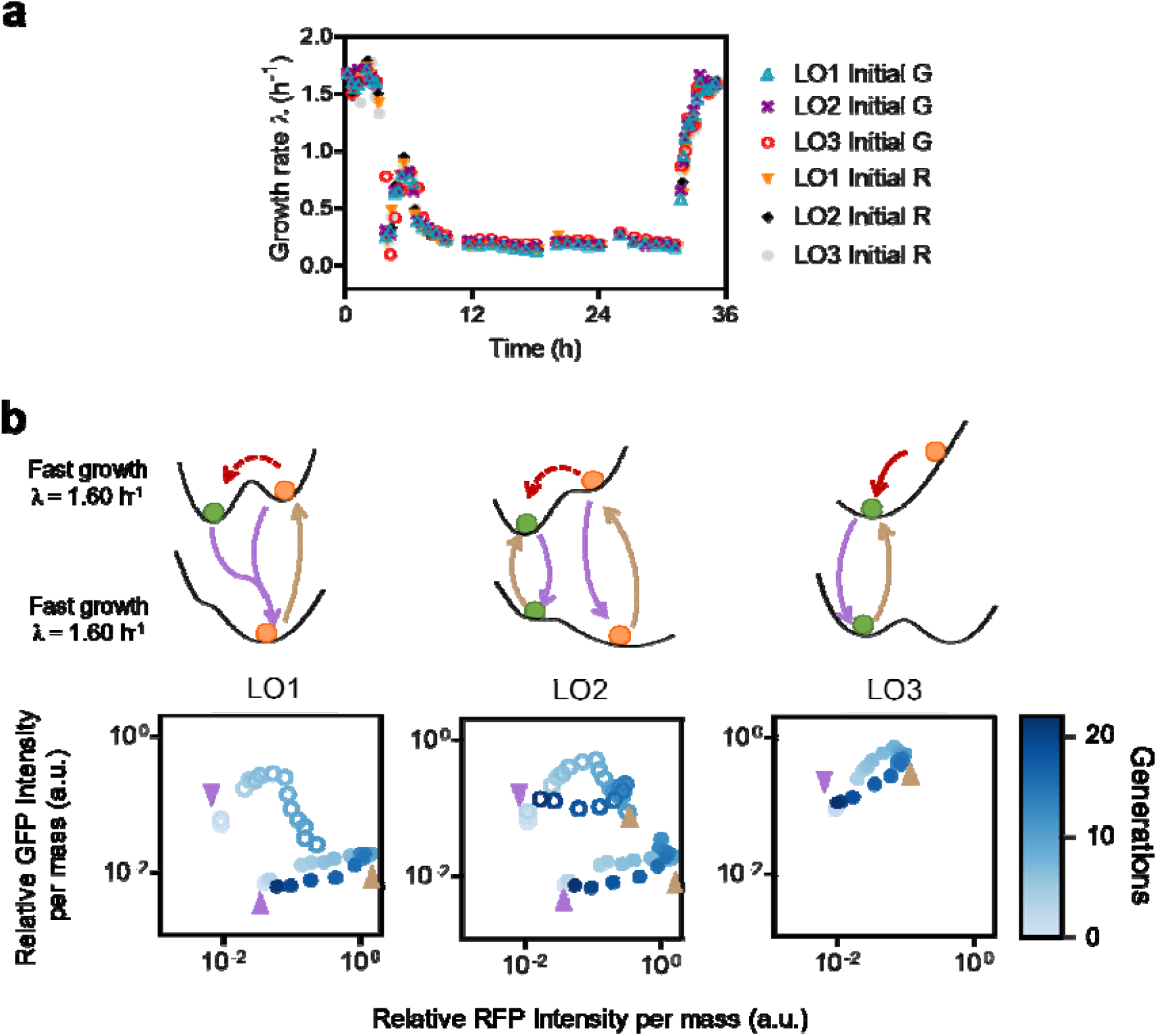
Cell fate determination trajectories during growth shift. **(a)**Time course of the instantaneous growth rate of cell cultures measured throughout nutrient downshift and upshift in batch culture. Nutrient shift experiments were performed between RDM glucose medium containing high-quality nitrogen sources (mixture of amino acids and ammonia) and MOPS (-NH_4_Cl) glucose minimal medium supplemented with glutamate as the sole nitrogen source. The instantaneous growth rate is derived by calculating the derivative of OD_600_concerning time . The instantaneous growth rate dropped abruptly after the nutrient downshift. This can be explained by the growth adaptation right after the depletion of amino acids and ammonia when the metabolic bottleneck for amino acid synthesis is significant. Rapid recovery and overshot its post-shift steady-state value until 0.9 h^-1^was observed, presumably because of the relatively high level of glucose uptake. The subsequent slowing down was observed until the final exponential growth rate at approximately 0.18 h^-1^. Growth rate data from three different designs of circuits (LO1, LO2, and LO3) triggered to two distinct initial states (G and R) are plotted. These data are quite repeatable, indicating that the cell growth dynamics are not affected by different genetic circuits. **(b)**Growth downshift and upshift experiments were carried out with cells initially triggered to the red state (pre-cultured in RDM glucose medium supplied with 0.2 mM IPTG, see **Methods**). Noise-induced spontaneous state transitions from red state to green state under fast growth conditions were identified after removing the inducer. For strains LO1 and LO2, only a small proportion of cells flip (red dashed lines). This observation is consistent with the result shown in **Extended Data Fig.10d**. In the case of LO3, all red state cells were destabilized, leading to the green state under fast growth (red solid line). The high RFP state is maintained throughout the growth downshift as well as the subsequent upshift for LO1 and LO2 strains.

**Extended Data Fig. 9:**
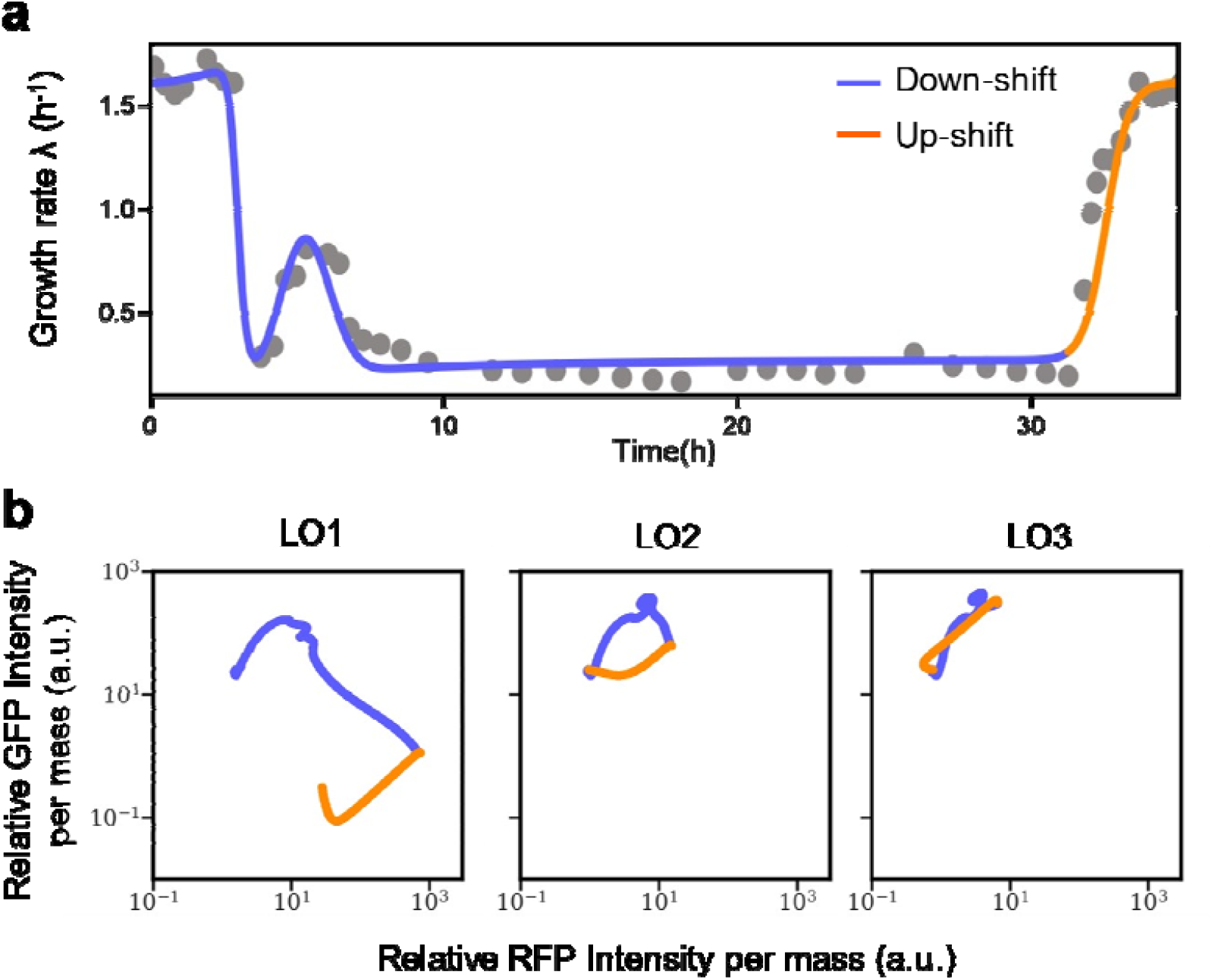
Cell fate determination trajectories of initial green state cells predicted by simulation. **(a)**Curve fitting of instantaneous growth rate across nutrient upshift and downshift conditions. The grey scatters are the instantaneous growth rate, and the coloured solid line denotes the growth rate which is fitted using a series of Hill functions. The slate blue part indicates the nutrient downshift, and the orange one shows the nutrient-upshift condition. **(b)**We applied the deterministic model (**Supplementary Notes 1**and **3**) to capture the growth-mediated dynamics of cell fate decision trajectories for each of the systems (**Fig. 4b**). The dynamics of model prediction closely follow the main features of the population-averaged dynamics of cell fate determination for each case. Three panels plot the predicted cell-fate-decision trajectories of strains LO1, LO2, and LO3, respectively.

**Extended Data Fig. 10:**
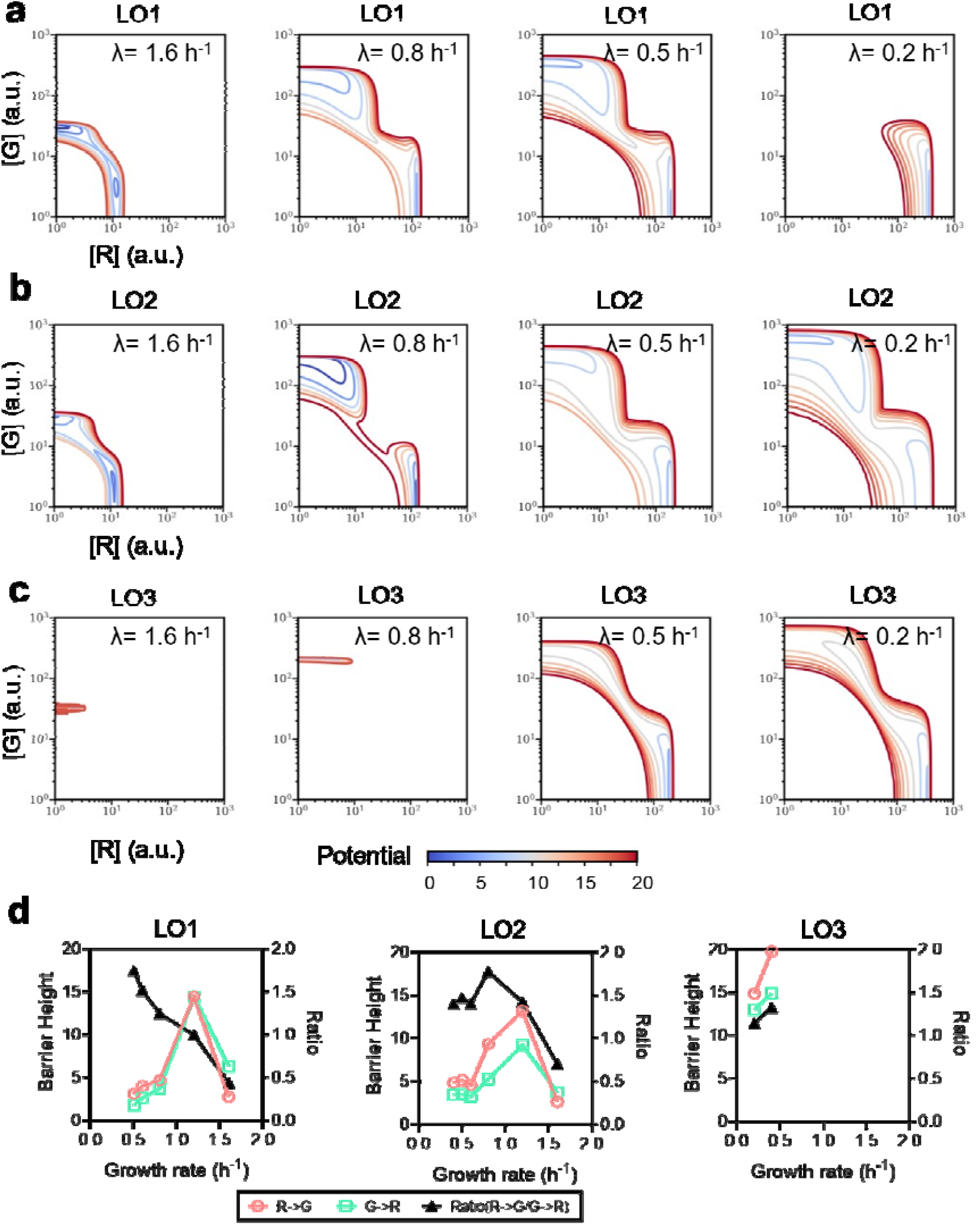
Probability potential landscape is reshaped during growth-rate fluctuation. Computed potential landscapes of the mutual repression switch illustrate the robustness of the states of cells with different growth rates and genetic backgrounds (**Supplementary Note 5**). The contour lines indicate the potential at the specific protein concentration ([G] and [R] axes). **(a-c)**Strains growing at varied growth rates show distinct potential landscapes. **(a)**As the decreasing of the growth rate, the bistability of the strain LO1 remained when the growth rate is greater than the critical growth rate, however, the bistability catastrophe happens at the critical growth rate, and the system remains the red stable state only when the growth rate is lower than the critical growth rate. **(b)**The strain LO2 remaining two stable states in our experimental conditions, the computed potential landscapes indicate the intrinsic parameters of promoter make the system behave more robustly to the fluctuation in growth rate. **(c)**The strain LO3 shows bistability only at slow growth rates. **(d)**the relation between growth rate and barrier height of the strains LO1, LO2, and LO3. The green squares denote the barrier heights between the G state and saddle point (G→R), and the red circles indicate the barrier heights between the R state and saddle point (R→G). The black triangles denote the relative changes of the two barrier heights (R→G/G→R).

## References

1 Balázsi, G., van Oudenaarden, A.& Collins, J. J.Cellular decision making and biological noise: from microbes to mammals. Cell144, 910–925, (2011).

2 Slack, J. M.& Dale, L.Essential developmental biology. (John Wiley & Sons, 2021).

3 Waddington, C.The Strategy of the Genes: A Discussion of Some Aspects of Theoretical Biology. (London: *λ*llen and Unwin, 1957).

4 Hanna, J.et al.Direct cell reprogramming is a stochastic process amenable to acceleration. Nature462, 595–601, (2009).

5 Ahrends, R.et al.Controlling low rates of cell differentiation through noise and ultrahigh feedback. Science344, 1384–1389, (2014).

6 Davidson, E. H.The regulatory genome: gene regulatory networks in development and evolution. (Elsevier, 2010).

7 Peter, I. S.& Davidson, E. H.Assessing regulatory information in developmental gene regulatory networks. Proc. Natl. Acad. Sci. USA. 114, 5862–5869, (2017).

8 Zaret, K. S.Pioneer transcription factors initiating gene network changes. Annual review of genetics54, 367, (2020).

9 Zhou, J. X.& Huang, S.Understanding gene circuits at cell-fate branch points for rational cell reprogramming. Trends Genet. 27, 55–62, (2011).

10 Iglesias-Bartolome, R.& Gutkind, J. S.Signaling circuitries controlling stem cell fate: to be or not to be. Current opinion in cell biology23, 716–723, (2011).

11 Kussell, E.& Leibler, S.Phenotypic diversity, population growth, and information in fluctuating environments. Science309, 2075–2078, (2005).

12 Arkin, A., Ross, J.& McAdams, H. H.Stochastic kinetic analysis of developmental pathway bifurcation in phage A-infected Escherichia coli cells. Genetics149, 1633–1648, (1998).

13 Oppenheim, A. B.et al.Switches in bacteriophage lambda development. Annual review of genetics39, 409–429, (2005).

14 Fujita, M.& Losick, R.Evidence that entry into sporulation in Bacillus subtilis is governed by a gradual increase in the level and activity of the master regulator Spo0A. Genes & development19, 2236–2244, (2005).

15 Vlamakis, H., Aguilar, C., Losick, R.& Kolter, R.Control of cell fate by the formation of an architecturally complex bacterial community. Genes & development22, 945–953, (2008).

16 Pastushenko, I.et al.Identification of the tumour transition states occurring during EMT. Nature556, 463–468, (2018).

17 McFaline-Figueroa, J. L.et al.A pooled single-cell genetic screen identifies regulatory checkpoints in the continuum of the epithelial-to-mesenchymal transition. Nat. Genet. 51, 1389–1398, (2019).

18 Pastushenko, I.& Blanpain, C.EMT transition states during tumor progression and metastasis. Trends Cell Biol. 29, 212–226, (2019).

19 Gardner, T. S., Cantor, C. R.& Collins, J. J.Construction of a genetic toggle switch in Escherichia coli. Nature403, 339–342, (2000).

20 Litcofsky, K. D.et al.Iterative plug-and-play methodology for constructing and modifying synthetic gene networks. Nat. Methods9, 1077–1080, (2012).

21 Cameron, D. E.& Collins, J. J.Tunable protein degradation in bacteria. Nat. Biotechnol. 32, 1276–1281, (2014).

22 Klumpp, S.& Hwa, T.Growth-rate-dependent partitioning of RNA polymerases in bacteria. Proc. Natl. Acad. Sci. USA;. 105, 20245–20250, (2008).

23 Klumpp, S., Zhang, Z.& Hwa, T.Growth rate-dependent global effects on gene expression in bacteria. Cell139, 1366–1375, (2009).

24 Scott, M.et al.Interdependence of cell growth and gene expression: origins and consequences. Science330, 1099–1102, (2010).

25 You, C.et al.Coordination of bacterial proteome with metabolism by cyclic AMP signalling. Nature500, 301–306, (2013).

26 Dai, X.et al.Reduction of translating ribosomes enables Escherichia coli to maintain elongation rates during slow growth. Nat. Microbiol. 2, 16231, (2016).

27 Erickson, D. W.et al.A global resource allocation strategy governs growth transition kinetics of Escherichia coli. Nature551, 119–123, (2017).

28 Basan, M.et al.A universal trade-off between growth and lag in fluctuating environments. Nature584, 470–474, (2020).

29 Zhang, R.et al.Topology-dependent interference of synthetic gene circuit function by growth feedback. Nat. Chem. Biol16, 695–701, (2020).

30 Tan, C., Marguet, P.& You, L.Emergent bistability by a growth-modulating positive feedback circuit. Nat. Chem. Biol5, 842–848, (2009).

31 Li, G.-W., Burkhardt, D., Gross, C.& Weissman, J. S.Quantifying absolute protein synthesis rates reveals principles underlying allocation of cellular resources. Cell157, 624–635, (2014).

32 Strogatz, S. H.Nonlinear dynamics and chaos: with applications to physics, biology, chemistry, and engineering. (CRC press, 2018).

33 Bhattacharya, S., Zhang, Q.& Andersen, M. E.A deterministic map of Waddington’s epigenetic landscape for cell fate specification. BMC Syst. Biol. 5, 1–12, (2011).

34 Wu, M.et al.Engineering of regulated stochastic cell fate determination. Proc. Natl. Acad. Sci. USA110, 10610–10615, (2013).

35 Moris, N., Pina, C.& Arias, A. M.Transition states and cell fate decisions in epigenetic landscapes. Nat. Rev. Genet. 17, 693–703, (2016).

36 Oehler, S., Eismann, E. R., Krämer, H.& Müller‐Hill, B.The three operators of the lac operon cooperate in repression. EMBO J. 9, 973–979, (1990).

37 Soldatov, R.et al.Spatiotemporal structure of cell fate decisions in murine neural crest. Science364, (2019).

38 Coomer, M. A., Ham, L.& Stumpf, M. P.H. Noise distorts the epigenetic landscape and shapes cell-fate decisions. Cell Syst., (2021).

39 Aoki, S. K.et al.A universal biomolecular integral feedback controller for robust perfect adaptation. Nature570, 533–537, (2019).

40 Li, Y.et al.A programmable fate decision landscape underlies single-cell aging in yeast. Science369, 325–329, (2020).

41 Bakshi, S.et al.Tracking bacterial lineages in complex and dynamic environments with applications for growth control and persistence. Nat. Microbiol. 6, 783–791, (2021).

42 Wang, J., Xu, L.& Wang, E.Potential landscape and flux framework of nonequilibrium networks: robustness, dissipation, and coherence of biochemical oscillations. Proc. Natl. Acad. Sci. USA. 105, 12271–12276, (2008).

43 Wang, J., Zhang, K., Xu, L.& Wang, E.Quantifying the Waddington landscape and biological paths for development and differentiation. Proc. Natl. Acad. Sci. USA108, 8257–8262, (2011).

44 Kang, X., Wang, J.& Li, C.Exposing the underlying relationship of cancer metastasis to metabolism and epithelial-mesenchymal transitions. Iscience21, 754–772, (2019).

45 Li, P.& Elowitz, M. B.Communication codes in developmental signaling pathways. Development146, dev170977, (2019).

46 Johnson, H. E.& Toettcher, J. E.Signaling dynamics control cell fate in the early Drosophila embryo. Dev. Cell48, 361-370. e363, (2019).

47 Vlamakis, H.et al.Sticking together: building a biofilm the Bacillus subtilis way. Nat. Rev. Microbiol. 11, 157–168, (2013).

48 García-Jiménez, C.& Goding, C. R.Starvation and pseudo-starvation as drivers of cancer metastasis through translation reprogramming. Cell Metab. 29, 254–267, (2019).

49 Schlissel, G.& Li, P.Synthetic developmental biology: understanding through reconstitution. Annu. Rev. Cell Dev. Biol. 36, 339–357, (2020).

50 Zorzan, I.et al.Synthetic designs regulating cellular transitions: Fine-tuning of switches and oscillators. Curr. Opin. Syst. Biol. 25, 11–26, (2021).

51 Shakiba, N., Jones, R. D., Weiss, R.& Del Vecchio, D.Context-aware synthetic biology by controller design: Engineering the mammalian cell. Cell Syst. 12, 561–592, (2021).

52 Negrete, J., Jr. & Oates, A. C.Towards a physical understanding of developmental patterning. Nat. Rev. Genet. 22, 518–531, (2021).

## References

53 Datta, S., Costantino, N.& Court, D. L.A set of recombineering plasmids for gram-negative bacteria. Gene379, 109–115, (2006).

54 Engler, C., Kandzia, R.& Marillonnet, S.A one pot, one step, precision cloning method with high throughput capability. PloS one3, e3647, (2008).

55 Neidhardt, F. C., Bloch, P. L.& Smith, D. F.Culture medium for enterobacteria. J. Bacteriol. 119, 736–747, (1974).

56 Zheng, H.et al.General quantitative relations linking cell growth and the cell cycle in Escherichia coli. Nat. Microbiol. 5, 995–1001, (2020).

57 Meyer, A. J.et al.Escherichia coli “Marionette” strains with 12 highly optimized small-molecule sensors. Nat. Chem. Biol15, 196–204, (2019).

58 Bren, A.et al.Glucose becomes one of the worst carbon sources for E.coli on poor nitrogen sources due to suboptimal levels of cAMP. Sci. Rep. 6, 24834, (2016).

59 Martin, M.Cutadapt removes adapter sequences from high-throughput sequencing reads. EMBnet j. 17, 10–12, (2011).

60 Langmead, B.& Salzberg, S. L.Fast gapped-read alignment with Bowtie 2. Nat. Methods9, 357–359, (2012).

61 Danecek, P.et al.Twelve years of SAMtools and BCFtools. Gigascience10, giab008, (2021).

62 Borujeni, A. E.et al.Genetic circuit characterization by inferring RNA polymerase movement and ribosome usage. Nat. Commun. 11, 1–18, (2020).

## References

22 Klumpp, S.& Hwa, T.Growth-rate-dependent partitioning of RNA polymerases in bacteria. Proc. Natl. Acad. Sci. USA. 105, 20245–20250, (2008).

63 Klumpp, S.Growth-rate dependence reveals design principles of plasmid copy number control. PLoS One6, e20403, (2011).

64 Zong, Y.et al.Insulated transcriptional elements enable precise design of genetic circuits. Nat. Commun. 8, 52, (2017).

