## supporting information for "Unbalanced response to growth variations reshapes the cell fate decision landscape"

**Affiliations:**

This section provides details of the model introduced in the main text. To explore the mutual repression circuit properties, including steady state, bistability, and phase diagrams, we first construct ODEs that describe the regulation relationship between two repressors. Analyzing the ODEs and correspondently dimensionless forms, insights into the origins of the bistability and hysteresis, and exhibits the bistability collapsing when the cells grow at different growth rates. For dissecting the underlying mechanism of the growth-rate-dependent gene expression, we propose a deterministic model that involves gene expression fluxes flowing in the central dogma. The dynamics of the circuit that undergo the nutrient up-and-down shift are simulated by a flux-controlled regulation model (FCR model) framework which was proposed by Erickson et al.^1^ Based on the FCR model, we apply the quasi-steady-state approximation to the biological processes whose timescales are separable from protein synthesis and capture the main features of the gene expression dynamics observed in experiments.

**Supplementary Note1: A deterministic model for the mutual repressive system**

To determine whether the cellular physiology or cell growth rate could account for the cell fate determination, we set up a simple mechanistic model (as illustrated in Extended Data Fig. 5) to describe and further predict the effect of cell growth rate on the cell fate determination of the mutual repression system:

$$\frac{d\left[ R \right]}{dt}=\alpha_{R}\cdot H_{R}\left( \left[ G \right] \right)-\lambda\cdot\left[ R \right]$$

(S1)

$$\frac{d\left[ G \right]}{dt}=\alpha_{G}\cdot H_{G}\left( \left[ R \right] \right)-\lambda\cdot\left[ G \right]$$

(S2)

where $\alpha$ refers to the protein synthesis rate, $\left[ R \right]$ and $\left[ G \right]$are the concentrations of TetR and LacI, respectively. The mutual inhibition was expressed by two Hill functions $H_{R}\left( \left[ G \right] \right)$ and $H_{G}\left( \left[ R \right] \right)$, which can be well approximated to the cooperative regulatory process. The protein decay is mainly due to the growth dilution $\lambda$ under the assumption that protein turnover is negligible. Hill function can be expressed as follows:

$$H_{R}\left( \left[ G \right] \right)=\tau_{R}+\frac{1-\tau_{R}}{1+\left( \frac{\left[ G \right]}{K_{DG}} \right)^{n_{G}}}$$

(S3)

$$H_{G}\left( \left[ R \right] \right)=\tau_{G}+\frac{1-\tau_{G}}{1+\left( \frac{\left[ R \right]}{K_{DR}} \right)^{n_{R}}}$$

(S4)

where $\tau$, $K$, and $n$ represent the promoter leakage ratio, dissociation constant (i.e., the repression threshold), and Hill coefficient. Unless otherwise mentioned, the theoretical values of the Hill coefficient, where $n_{G}=4$ and $n_{R}=2$, are used in the model.

Let us consider $\tilde{\alpha}_{G}=\alpha_{G}/\lambda$, $\tilde{\alpha}_{R}=\alpha_{R}/\lambda$, $dg=\lambda dt$, where the dimensionless $g$ represents the generation time of cells. Using the new parameters, the dimensionless equations can be written as:

$$\frac{d\left[ R \right]}{dg}=\tilde{\alpha}_{R}H_{R}\left( \left[ G \right] \right)-\left[ R \right]$$

(S5)

$$\frac{d\left[ G \right]}{dg}=\tilde{\alpha}_{G}H_{G}\left( \left[ R \right] \right)-\left[ G \right]$$

(S6)

Similarly, we introduce dimensionless protein input terms, $\tilde{H}_{R}$ and $\tilde{H}_{G}$ are:

$$\tilde{H}_{R}\left( \left[ \tilde{G} \right] \right)=\tau_{R}+\frac{1-\tau_{R}}{1+\left( \left[ \tilde{G} \right]/\tilde{K}_{DG} \right)^{n_{G}}}$$

(S7)

$$\tilde{H}_{G}\left( \left[ \tilde{R} \right] \right)=\tau_{G}+\frac{1-\tau_{G}}{1+\left( \left[ \tilde{R} \right]/\tilde{K}_{DR} \right)^{n_{R}}}$$

(S8)

where $\left[ \tilde{G} \right]=\left[ G \right]/\tilde{\alpha}_{G}$, $\left[ \tilde{R} \right]=\left[ R \right]/\tilde{\alpha}_{R}$, $\tilde{K}_{DG}=K_{DG}/\tilde{\alpha}_{G}$ and $\tilde{K}_{DR}=K_{DR}/\tilde{\alpha}_{R}$.

Notice that $\tilde{\alpha}$ refers to the steady protein concentration (i.e., protein expression capacity), which indicates the maximal level when the protein expression is unregulated, the $\left[ \tilde{R} \right]$ and $\left[ \tilde{G} \right]$ can be regarded as the expression ratio of the steady state expression concentration over specific growth conditions.

In our system, $\tilde{\alpha}_{R}$ and $\tilde{\alpha}_{G}$ were determined experimentally by measuring the steady-state fluorescence intensity per mass of the reporter under different growth conditions of the two constitutively expressed parts (Fig. 2a, Extended Data Fig. 4a and 4b). Because of the strong growth rate dependence on protein expression level, our model introduces the dimensionless forms of protein concentration $\left[ \tilde{R} \right]$ and $\left[ \tilde{G} \right]$, which are normalized to the 0-1 range. $\tilde{K}_{DG}$ and $\tilde{K}_{DR}$ are the dimensionless repression thresholds of LacI and TetR, respectively, to their corresponding binding sites, in the unit of the expression capacity under different conditions.

Thus, we can rewrite the ODEs as follows:

$$\frac{\text{d}\left[ \tilde{R} \right]}{\text{d}g}=\tilde{H}_{G}\left( \left[ \tilde{G} \right] \right)-\left[ \tilde{R} \right]$$

(S9)

$$\frac{\text{d}\left[ \tilde{G} \right]}{\text{d}g}=\tilde{H}_{R}\left( \left[ \tilde{R} \right] \right)-\left[ \tilde{G} \right]$$

(S10)

The nullclines in Fig. 2b are drawn based on the abovementioned dimensionless model. Evaluating $\text{d}\left[ \tilde{R} \right]/\text{d}g=0$ and $\text{d}\left[ \tilde{G} \right]/\text{d}g=0$, we can get the stable steady states, $\left[ \tilde{R} \right]^{*}$ and $\left[ \tilde{G} \right]^{*}$, as well as the unstable one (saddle point) if exists. In this model, $\tilde{K}_{DG}$ and $\tilde{K}_{DR}$, which are growth rate dependent, are key parameters that determine the stability of the system. Fig. 2c is shown the growth-rate dependency of the dimensionless repression thresholds.

**Supplementary Note 2: Model of constitutive gene expression as a function of growth rate**

Our experimental data (Fig. 2a and Extended Data Fig. 4) have shown that a constitutive expression gene has varied protein synthesis rates by tuning the nutrient components of growth media. Furthermore, the response of synthesis rates of different proteins to growth rate variations is always unbalanced. In this section, we provide a deviation of the growth rate dependence feature of protein synthesis rate, using some empirical relations and key assumptions of bacterial “growth law” developed by Terence Hwa’s lab, as well as the RNA sequencing data from this work.

Here, we list the key assumptions underpinning the formulation of the model and interpret the biological meaning of each parameter.

Transcription modulation

The biological processes from DNA to mRNAs include DNA replication, mRNA synthesis, and their degradation. The overlap of DNA replication rounds^2^ and plasmid copy number controlling mechanism^3^ results in the gene copy number variation when varying the growth rate. The limited RNA polymerase pool regulates the transcription initial frequency^4^. The mRNA contents and the translation rate difference give rise to the variable lifetime of mRNAs^5^. These processes are still elusive and need to be carefully evaluated. Thus, instead of deducing a bottom-up model from the bioprocesses that include the details of DNA replication and transcription, we begin with the protein synthesis processes to model the protein synthesis rate and protein expression level.

With the aid of next-generation sequencing, we can determine the relative mRNA fraction of each gene (see Methods). Here, we introduce the mRNA abundance of a specific gene $i$, $\phi_{i}^{m}$:

$$\phi_{i}^{m}=\frac{m_{i}}{\sum m_{i}}$$

(S11)

where $m_{i}$ denotes the mRNA copy number of gene $i$, and $\sum m_{i}$ denotes the total mRNA copy number.

As shown in Extended Data Fig. 4d, the steady-state mRNA abundance of *tetR* and *lacI* genes show strong growth rate dependence. The relation between mRNA abundance and the growth rate of each gene was obtained by parameter fitting.

The protein translation rate is a function of the growth rate

Considering the simple model of gene expression, we assume gene $i$ has a synthesis rate $\alpha_{i}$, and its product concentration is $\left[ p_{i} \right]$. Protein only decays by cell growth dilution ($\lambda$). We have thus:

$$\frac{d\left[ p_{i} \right]}{dt}=\alpha_{i}-\lambda\left[ p_{i} \right]$$

(S12)

Here, the problem falls in the form of $\alpha_{i}$. Different from deducing the protein synthesis rate that depends on the specific concentration of gene expression machinery (S14), we propose a view that the protein synthesis rate $\alpha_{i}$ depends on cellular resource allocation strategy (Eq. (S17). It means that cells need to assign their limited resource to adapt to different nutrient conditions. Empirical growth laws give the quantitative relationship between growth rate and total protein $p$ translation:

$$\frac{d\left[ p \right]}{dt}=\kappa_{t}\left[ r_{a} \right]-\lambda\left[ p \right]$$

(S13)

where $\left[ p \right]$, $\kappa_{t}$, and $\left[ r_{a} \right]$ are total protein concentration, average ribosome translation rate, and concentration of active ribosome, respectively. Due to the integral feedback regulation mechanism of cell physiology^6^, the correlation between the protein synthesis rate $\alpha$ and steady-state growth rate $\lambda^{*}$ is well characterized experimentally and theoretically^6-8^.

The stoichiometric reaction form of protein expression can be given by

$$\frac{d\left[ p_{i} \right]}{dt}=k_{i}\left[ m_{i} \right]\left[ r_{a} \right]-\lambda\left[ p_{i} \right]$$

(S14)

where $k_{i}$ and $\left[ m_{i} \right]$ denote the translation initiation rate and the concentration of mRNA for gene $i$. Lump up all translation processes, and $\phi_{i}^{m}=\left[ m_{i} \right]/\left[ m \right]$, we have

$$\frac{\text{d}\left[ p \right]}{\text{d}t}=\frac{\text{d}\sum p_{i}}{\text{d}t}=\sum_{i} k_{i}\phi_{i}^{m}\left[ m \right]\left[ r_{a} \right]-\lambda\left[ p \right]$$

(S15)

where $\left[ m \right]$ is the concentration of total mRNA. When cells stay in a steady state, the protein components are relatively constant, we have $\text{d}\left[ p \right]/\text{d}t=0$ and $\text{d}\left[ p_{i} \right]/\text{d}t=0$. Note that the number fraction of protein $i$ is $\phi_{i}^{p}$, we introduce a parameter, translation efficiency $\zeta_{i}$ which can be deduced from Eqs. (S14) and (S15),

$$\zeta_{i}=\frac{\phi_{i}^{p}}{\phi_{i}^{m}}=\frac{k_{i}}{\sum_{i} k_{i}\phi_{i}^{m}}$$

(S16)

We suppose the translation efficiency for gene $i$ of the circuit, $\zeta_{i}$, are constants depending on RBS strength rather than the growth conditions. Revisiting the protein translation process, ribosomes bind to the 5' UTR region of an mRNA and initial the protein translation. The initial rate is strongly correlated to the ribosome binding site (RBS). Although post-transcriptional regulations, such as aptamers and miRNAs, may alter the translation initiation processes frequency depending on the specific environmental cues, they are not ubiquitous for most genes. Moreover, a recent study reveals that the translation efficiency of most genes is independent of growth conditions^9^. Therefore, comparing Eqs. (S13) and (S15), we have $\sum_{i} k_{i}\phi_{i}^{m}\left[ m \right]=\kappa_{t}$, combined with Eq. (S14), we have gene $i$ expression rate,

$$\frac{\text{d}\left[ p_{i} \right]}{\text{d}t}=\zeta_{i}\phi_{i}^{m}\kappa_{t}\left[ r_{a} \right]-\lambda\left[ p_{i} \right]$$

(S17)

Compare Eqs. (S12) and (S17), we have thus $\alpha_{i}=\zeta_{i}\phi_{i}^{m}\kappa_{t}\left[ r_{a} \right]$. In this study, we characterized the $\phi_{i}^{m}$ of the two TFs of the mutual repression circuit in various growth condition via RNA-seq (Extended Data Fig. 4d). Data were fitted by Hill functions for obtaining a smooth curve. General translation rates $\kappa_{t}$ and concentration of active ribosome $\left[ r_{a} \right]$ were collected from ref.^1^. For predicting the steady states of gene expression, the range of growth rates $\lambda$ that cross our experimental conditions are used, and $\alpha_{R,G}\left( \lambda\right)$ are obtained (Extended Data Fig. 4g). Parameters are given in Supplementary Table 9.

**Supplementary Note 3: Model for gene expression in fluctuated-growth condition.**

The model mentioned above described the mechanism of the growth-rate-dependent gene expression which was raised by the global transcription regulation and ribosomal context. For predicting the gene expression and cellular growth dynamics in a fluctuated environment, Erikson et.al^1^ have dissected the process from nutrient influx to the protein synthesis flux and proposed a model frame, the flux-controlled regulation (FCR) model. In our study, we perturbated cells’ growth rates by up-shift and down-shift nutrient quality of growth media and characterized the growth rates and the circuit dynamics (Fig. 4 and Extended Data Fig. 8).

We supposed that biomass accumulation is the origin of the protein synthesis, and the accumulation rate $\lambda$ (i.e., growth rate) indicates the total protein synthesis rate, $\alpha=\kappa_{t}\left[ r_{a} \right]$, which is contributed by two factors: average ribosome translation rate, and the concentration of active ribosomes. In the previous study from Erickson et al.^1^, a key parameter, the translational activity of the ribosomes $\sigma$, was introduced to connect the growth rate $\lambda$ and the gene expression rate $\alpha$.

The translational activity of the ribosomes $\sigma$ is the coarse-grained protein translation rate by assuming every ribosome contributes equally to protein synthesis, and its relationship with growth rate $\lambda$ can be given by:

$$M_{p}\cdot\lambda= \sigma R ,$$

(S18)

where $M_{p}$ represents total protein mass per standard volume, $R$ is the ribosomes and their affiliated factors in a standard volume, and we have

$$\sigma\left( t \right)=\lambda\left( t \right)/\phi_{R}\left( t \right),$$

(S19)

where $\phi_{R}\left( t \right)$ is the mass fraction of ribosomal proteins in total protein mass. Based on the empirical growth law^10^, the relationship between growth rate $\lambda$ and $\phi_{R}$ is given by a linear function:

$$\phi_{R}^{*}=\lambda^{*}/\kappa_{R}+\phi_{R,0},$$

(S20)

where $\phi_{R,0}$ denotes the basal level of $\phi_{R}$, when growth rate $\lambda$ equals to 0, and $\kappa_{R}$ is the maximal translation rate of the ribosome. For describing the protein synthesis flux allocated for ribosome synthesis, we start with the homeostasis of $\phi_{R}$,

$$\frac{\text{d}R}{\text{d}t}=\psi_{R}\sigma R$$

(S21)

where $\psi_{R}$ represents the fraction of total protein synthesis flux that is allocated to regenerate ribosomes and we call it the global regulatory function for ribosome synthesis.

$$\frac{\text{d}\phi_{R}}{\text{d}t}\equiv\frac{\text{d}}{\text{d}t}\left( \frac{R}{M} \right)=\frac{1}{M}\frac{\text{d}R}{\text{d}t}-\lambda\cdot\phi_{R}$$

(S22)

When the cells are in a steady state, all cellular contexts are relatively invariable, and we have $d\phi_{R}/dt=0$,

$$\psi_{R}^{*}=\frac{\lambda^{*}}{\sigma^{*}}$$

(S23)

According to Eq. (S19) we have $\psi_{R}^{*}=\phi_{R}^{*}$.

Back to the molecular mechanism underpinning the regulation of the translational activity, the available metabolites determine the magnitude of the translational activity in a fluctuating environment. We suppose the timescale of the regulation of the translational activity is much faster than gene expression regulation, using quasi-steady approximation, we have $\sigma\left( t \right)=\sigma^{*}$. We noticed that the global regulation function for a specific gene is also leaded by the available metabolites (the promotor activity of a specific gene is regulated by ppGpp or cAMP, and the levels of these signal molecules are determined by metabolites levels in cells), therefore, during the growth fluctuation, the effects that the available metabolites exert on the global regulatory function can be mimicked through the effects on translational activity, i.e, $\psi\left( t \right)\approx\hat{\psi}(\sigma(t))$. To sum up, we can have the global regulation function for ribosome expression,

$$\psi_{R}\approx\hat{\psi}_{R}\left( \sigma^{*} \right)=\phi_{R}^{*}\left( \lambda\left( \sigma^{*} \right) \right)=\frac{\phi_{R,0}}{1-\kappa_{R}\sigma^{*}},$$

(S24)

where $\lambda\left( \sigma^{*} \right)=\sigma^{*}\cdot\phi_{R}\left( t \right)$.

According to Eq. (S22), we have the dynamics of the $\phi_{R}\left( t \right)$,

$$\frac{\text{d}\phi_{R}}{\text{d}t}=\frac{\phi_{R,0}\sigma}{1-\kappa_{R}\sigma}\phi_{R}-\lambda\phi_{R}$$

(S25)

The abovementioned equation describes how translational activity affects the dynamics of ribosome contents. Next, we are going to discuss the synthesis flux allocated to synthesise the circuit protein. When the cells stay in a steady state, we have

$$\left[ p_{i} \right]^{*}=\frac{\alpha^{*}}{\lambda^{*}}$$

(S26)

However, Eq. (S26) cannot describe the protein levels when the cellular contexts are fluctuating in a perturbating niche. To obtain the general dynamics of the genetic circuits’ gene expression, considering the expression dynamics of the circuit gene $i$ and we have

$$\frac{\text{d}\left[ p_{i} \right]}{\text{d}t}=\psi_{i}\sigma\phi_{R}-\lambda\left[ p_{i} \right].$$

(S27)

When the system is in a steady state, we have $\psi_{i}^{*}=\left[ p_{i} \right]^{*}$. Considering the gene expression characterized by Eq. (S17) and $\psi\left( t \right)\approx\hat{\psi}\left( \sigma\left( t \right) \right)$, we have

$$\psi_{i}\left( t \right)\approx\hat{\psi}_{i}\left( t,\sigma^{*} \right)=\frac{\zeta_{i}\phi_{i}^{m}\left( \lambda\left( \sigma^{*} \right) \right)\cdot\kappa_{t}\left( \lambda\left( \sigma^{*} \right) \right)\cdot\left[ r_{a} \right]\left( \lambda\left( \sigma^{*} \right) \right)}{\lambda\left( \sigma^{*} \right)}.$$

(S28)

In our study, we performed the nutrient up- and down-shift experiment (Fig. 4 and Extended Data Fig. 8), the cells’ state is tuned to one state using an inducer (For LacI/GFP state, cTc is used; for TetR/RFP state, IPTG is used) at the seed culture stage. The cells are cultured at the initial medium for more than 10 generation time without an inducer and reach a steady state. At $t_{1}$, growth culture is diluted to a poor nutrient quality medium and cultured for another 10 generations. Finally, the growth culture is inoculated to high nutrient quality medium at $t_{2}$. We can obtain the empirical relationship between growth rate and time, $\lambda\left( t \right)$, by calculating the instantaneous growth rate.

According to the Eqs. (S1)(S2)(S19)(S25)(S27)(S28), we can evaluate the dynamics of the circuit in perturbated conditions by following equations,

$$\frac{\text{d}\phi_{R}}{\text{d}t}=\frac{\phi_{R,0}\sigma}{1-\sigma/\kappa_{R}}\phi_{R}-\lambda\phi_{R}$$

(S29)

$$\sigma\left( t \right)=\lambda\left( t \right)/\phi_{R}\left( t \right)$$

(S30)

$$\frac{\text{d}\left[ G \right]}{\text{d}t}=\hat{\psi_{G}}\left( \lambda\left( \sigma\right) \right)\sigma\phi_{R}H_{G}\left( \left[ R \right] \right)-\lambda\left[ G \right]$$

(S31)

$$\frac{\text{d}\left[ R \right]}{\text{d}t}=\hat{\psi_{R}}\left( \lambda\left( \sigma\right) \right)\sigma\phi_{R}H_{R}\left( \left[ G \right] \right)-\lambda\left[ R \right]$$

(S32)

where Eq.(S29) depicts the dynamics of $\phi_{R}\left( t \right)$, Eq. (S30) portrays the dynamics of $\sigma\left( t \right)$. In Eqs. (S31) and (S32), the term $\hat{\psi}\sigma\phi_{R}$ denotes the dynamics of the protein synthesis rate $\alpha$, and the terms $H_{G}\left( \left[ R \right] \right)$ and $H_{R}\left( \left[ G \right] \right)$ are Hill function terms which are identical to the terms in Eqs. (S1) and (S2).

To specify the initial values of the model, we calculate the steady-state solution of the mutual repressive system with the fixed growth rate at $t_{1}$, as well as the protein synthesis rates $\alpha_{R}^{*}\left( t_{1} \right)$ and $\alpha_{G}^{*}\left( t_{1} \right)$ (according to the empirical growth rate dependent relations given in Supplementary Table 9). The ribosome translation activity $\sigma^{*}\left( t_{1} \right)$ can be determined via Eq. (S30). Solving the fixed point (G state) of Eqs. (S1) and (S2) , we have $\left[ R \right]^{*}\left( t_{1} \right)$ and $\left[ G \right]^{*}\left( t_{1} \right)$ (Parameters are listed in Supplementary Table 9). The calculated cell fate determination trajectories are given in Extended Data Fig. 9. The dynamics of model prediction closely follow the main features of the population-averaged dynamics of cell fate determination shown in Fig. 4.

**Supplementary Note 4: A deterministic quasi-potential landscape**

Bhattacharya et al.^11^ proposed a simple method to map the quasi-potential landscape derived directly from the deterministic equations of the gene regulatory network. In the present work, we adopt this approach to visualize a one-dimensional quasi-potential landscape of our mutual repressive system, which provides a measure of the stability of the system.

A two-variable term $U$ is defined as the elevation of a quasi-potential along a given trajectory in a two-dimensional space. The local minima on this potential $U$ would correspond to the stable steady states of the system. Given that at the local minima, the rate of change in the expression of both $x$ and $y$ would vanish (i.e., $\partial U/\partial x=0$ and $\partial U/\partial y=0$). The change in quasi-potential $\Delta U$ can be given as follows:

$$\Delta U\left( x,y \right)=\frac{\partial U}{\partial x}\cdot\Delta x+\frac{\partial U}{\partial y}\cdot\Delta y$$

$$=-\frac{\text{d}x}{\text{d}t}\cdot\Delta x-\frac{\text{d}y}{\text{d}t}\cdot\Delta y$$

(S33)

where $\Delta x$ and $\Delta y$ are sufficiently small along the trajectory such that the vector force $\text{d}x/\text{d}t$ and $\text{d}y/\text{d}t$ that derive the system can be assumed to remain unchanged within the interval $\left[ \left( x,x+\Delta x \right);\left( y,y+\Delta y \right) \right]$.

In our case, to visualize the growth-rate dependent quasi-potential landscape of such a multi-dimensional system, we simplified the landscape to a one-dimensional one by substituting one variable with its steady-state solution^12^ (i.e., $\text{d}y/\text{d}t=0$):

$$\Delta U\left( x,y^{*} \right)=-\frac{\text{d}x}{\text{d}t}\cdot\Delta x$$

(S34)

One can obtain the overall change of the quasi-potential along a given trajectory by integrating $\Delta U$ of the system. We then map the one-dimensional quasi-potential landscape using our dimensionless model of the mutual repressive circuit:

$$U\left( x \right)=-\int_{0}^{x} \left( \frac{\text{d}\left[ \tilde{R} \right]}{\text{d}g} \right)\text{d}\left[ \tilde{R} \right]$$

(S35)

where $\text{d}\left[ \tilde{\text{R}} \right]/\text{d}g$ is given by Eq. (S8) and (S9) by taking the steady state solution of Eq. (S10), i.e.,

$$\left[ \tilde{G} \right]^{*}=\tau_{G}+\frac{1-\tau_{G}}{1+\left( \left[ \tilde{R} \right]/{\tilde{K}_{DR}} \right)^{n_{R}}}$$

(S36)

The initial value of the quasi-potential at the start of the trajectory where $\left[ \tilde{R} \right]=0$ is arbitrarily set to zero. The one-dimensional quasi-potential landscapes under various growth rates are given in Fig. 2d, Supplementary Fig. 3 and 4. This quasi-potential landscape provides a representation of the stability of cellular states under a given condition, for the local minima correspond to the stable states of a system. However, this represents neither escape time between states (i.e., state transition rate) nor the dynamics of developmental trajectories.

**Supplementary Note 5: A probability potential landscape**

Instead of the averaged deterministic dynamics (Extended Data Fig. 9) which might not capture some observations throughout the growth shift nor the spontaneous state transition under balanced growth, we consider a probabilistic description to model the cellular process of cell fate decision. The stochastic dynamics of the toggle switch can be described by the following Langevin equations:

$$\frac{\text{d}G}{\text{d}t}=\alpha_{G} H_{G}\left( \left[ R \right] \right)-\lambda\left[ G \right]+ \zeta_{G}$$

(S37)

$$\frac{\text{d}R}{\text{d}t}=\alpha_{R} H_{R}\left( \left[ G \right] \right)-\lambda\left[ R \right] +\zeta_{R}$$

(S38)

where $\zeta_{G}$ and $\zeta_{R}$ are two independent white Gaussian noise terms. In our study, we assumed that the noise is homogeneous. The covariance matrix is $\left\langle\zeta_{i}\left( t \right),\zeta_{j}\left( t' \right) \right\rangle=2\sqrt{D_{i}D_{j}}\delta_{i,j}\delta\left( t-t^{'} \right);i,j\in\{G,R\}$, where $\delta_{i,j}=0$ (if $i \neq j$), and $\delta_{i,j}=1$ (if $i=j$). The temporal change of the probability density function $P\left( G,R,t \right)$ is governed by a two-dimensional Fokker-Planck equation (FPE) (S39):

$$\frac{\partial P\left( G, R,t \right)}{\partial t}=-\frac{\partial}{\partial G}\left[ F_{1}\left( G, R \right)P \right]-\frac{\partial}{\partial R}\left[ F_{2}\left( G, R \right)P \right]$$

$$+\frac{\partial^{2}}{\partial G^{2}}\left[ D_{G} P \right]+\frac{\partial^{2}}{\partial R^{2}}\left[ D_{R} P \right].$$

(S39)

where the drift terms are the $F_{1}\left( G,R \right) = \alpha_{G}H_{G}\left( R \right)-\lambda G$ and $F_{2}\left( G,R \right) = \alpha_{R} H_{R}\left( G \right)-\lambda R$.

Wang et al.^13,14^ have proposed a theoretical framework, the potential landscape $U$, for evaluating the robustness and coherence of a biological system, which can be deduced form following equation (S40),

$$U\left( G,R \right)=-\ln\left[ P\left( G,R,t\to\infty\right) \right].$$

(S40)

For defining the parameters $D_{G}$ and $D_{R}$, one supposes that they become constant under specific growth conditions, whereas the experiments observed the magnitudes of noise increased with decreasing growth rate (Supplementary Fig. 5), we tuned its value by estimation from the experimental data.

To generate potential landscapes $U$, we calculated the asymptotic state of $P\left( x \right)$. Specifically, Neumann boundary condition was applied for conserving probability. $\mu\left( G,R \right)$ of the initial probability $P\left( G,R,t_{0} \right)$ was set at the saddle point, and the probability of each grid point was given by the two-dimensional Gaussian distributions Eq.(S41). All grid points used were set to 1024 $\times$ 1024. Alternating direction implicit algorithm was used to solve the FPE. The parameters used are listed in Supplementary Tables 9 and 10.

$$f\left( \mathbf{x} \right)=\frac{1}{\sqrt{(2\pi)^{k}|\Sigma|}}\text{exp}\left( -\frac{(\mathbf{x}-\boldsymbol{\mu})^{T}\Sigma^{-1}(\mathbf{x}-\boldsymbol{\mu})}{2} \right),$$

(S41)

where $x=\left( G,R \right)$ , $\Sigma=\left[ \begin{matrix} \sigma_{G}^{2} & \\ & \sigma_{R}^{2} \end{matrix} \right]$, and $k=2$.

**Supplementary Figures**

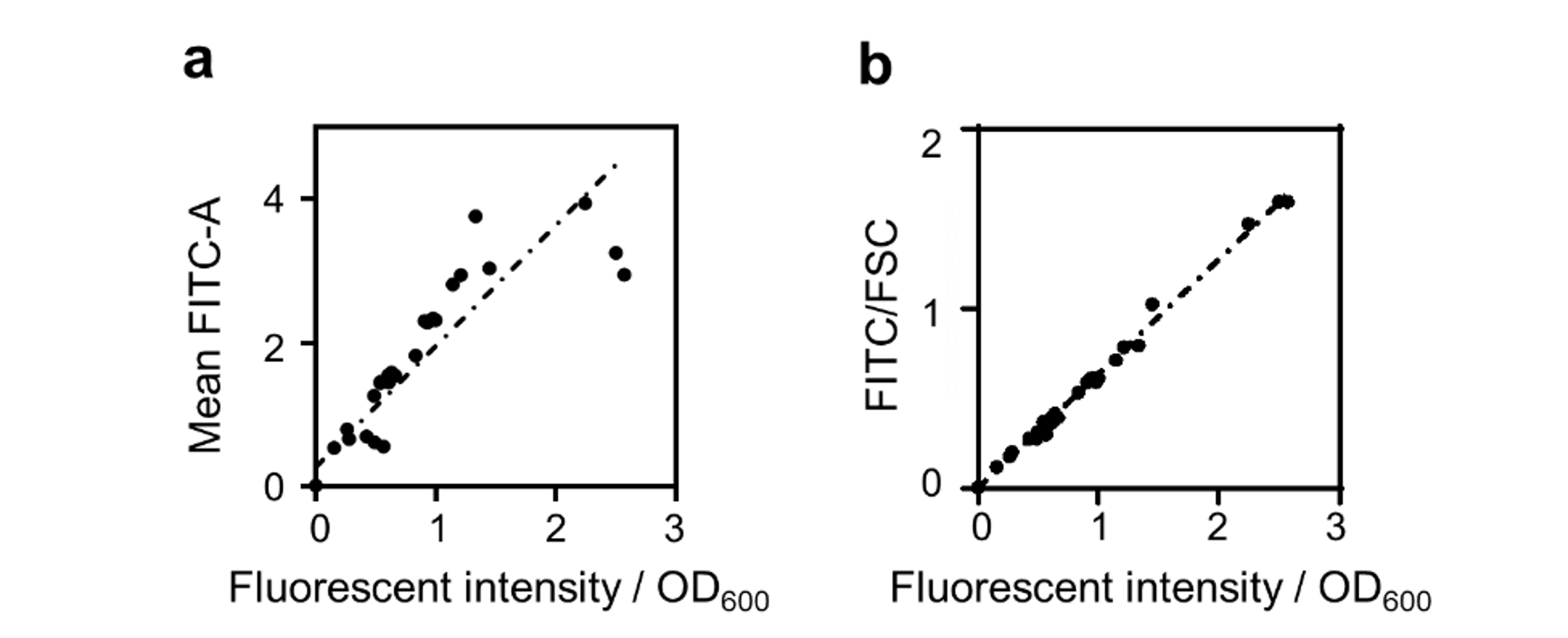

Supplementary Fig. 1: Validation of the method to quantify the fluorescent intensity per mass using flow cytometry. Constitutively expressed GFP using different promoters (i.e., *Ptrc2*, *P_L_tetO-1*, J23101) in combination with various RBSs (Ribosome binding sites) under diverse growth conditions (i.e., RDM glucose/glycerol, MOPS CAA glucose/glycerol, and RDM glucose with a set of sublethal doses of chloramphenicol) were provided. (a) The variation of cell size under different conditions could pose a limitation for flow cytometric analysis using the mean average value of the fluorescence signals (e.g., FITC-A), especially for quantitative analysis of gene expression over an extensive range of growth rates. For this purpose, FSC-H was used in this study to normalize the effect of cell size change in different growth conditions as its well linear relation with cell volume. (b) The linear relation (R^2^ > 0.99) between the FITC/FSC (ratio of mean FITC and mean FSC, see details in Materials and Methods) value obtained by flow cytometer and fluorescent intensity per OD_600_ obtained by a plate reader (microplate spectrophotometer, BioTek Epoch 2) is evident in numbers of constructs and conditions.

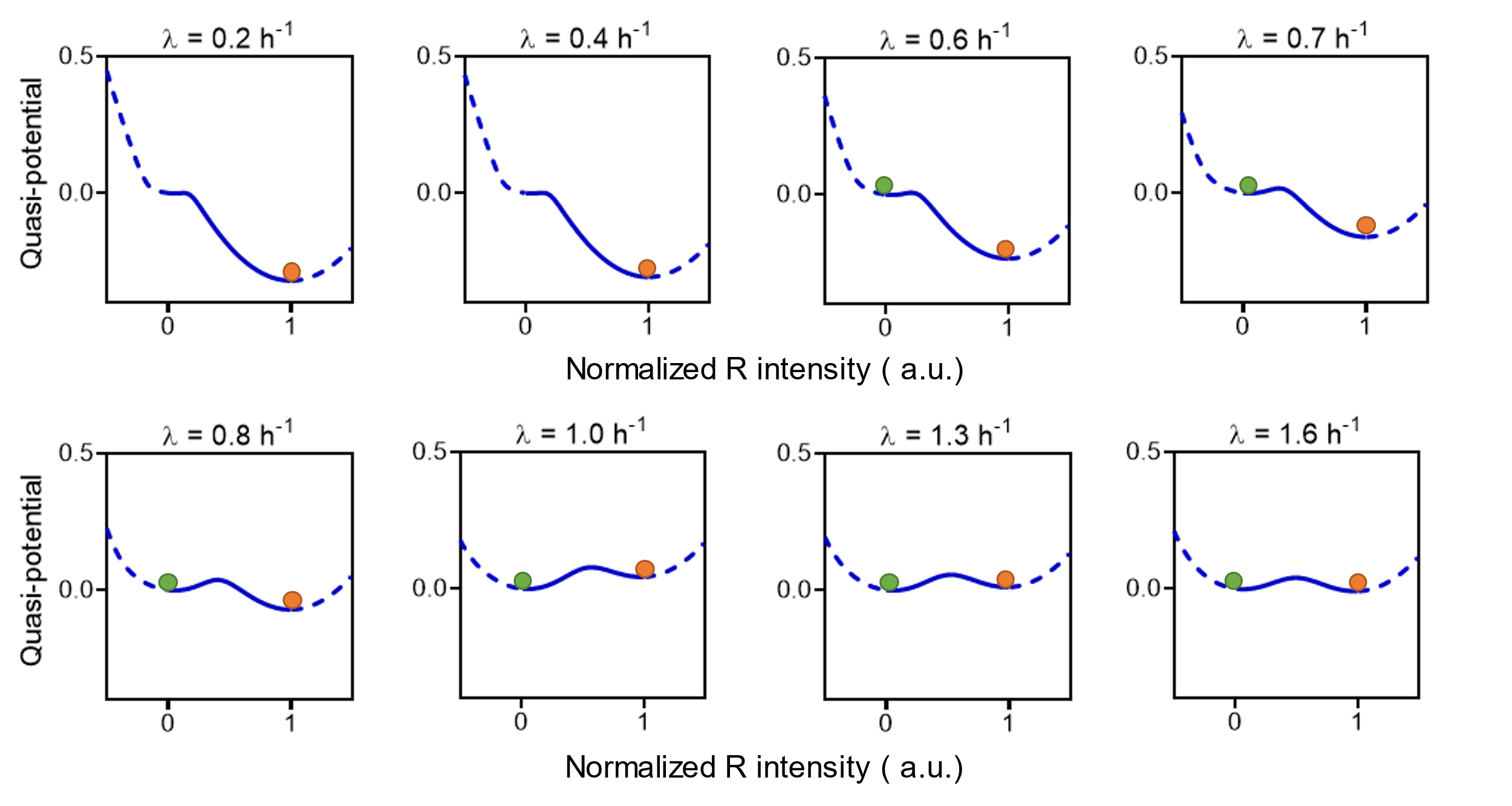

**Supplementary Fig. 2: Calculated one-dimensional quasi-potential landscapes for strain LO1 under selected growth rates.** The local minima and the top of the potential represent stable and unstable (saddle-point) fixed points, respectively. Stability is determined by the curvature at the stable fixed point.

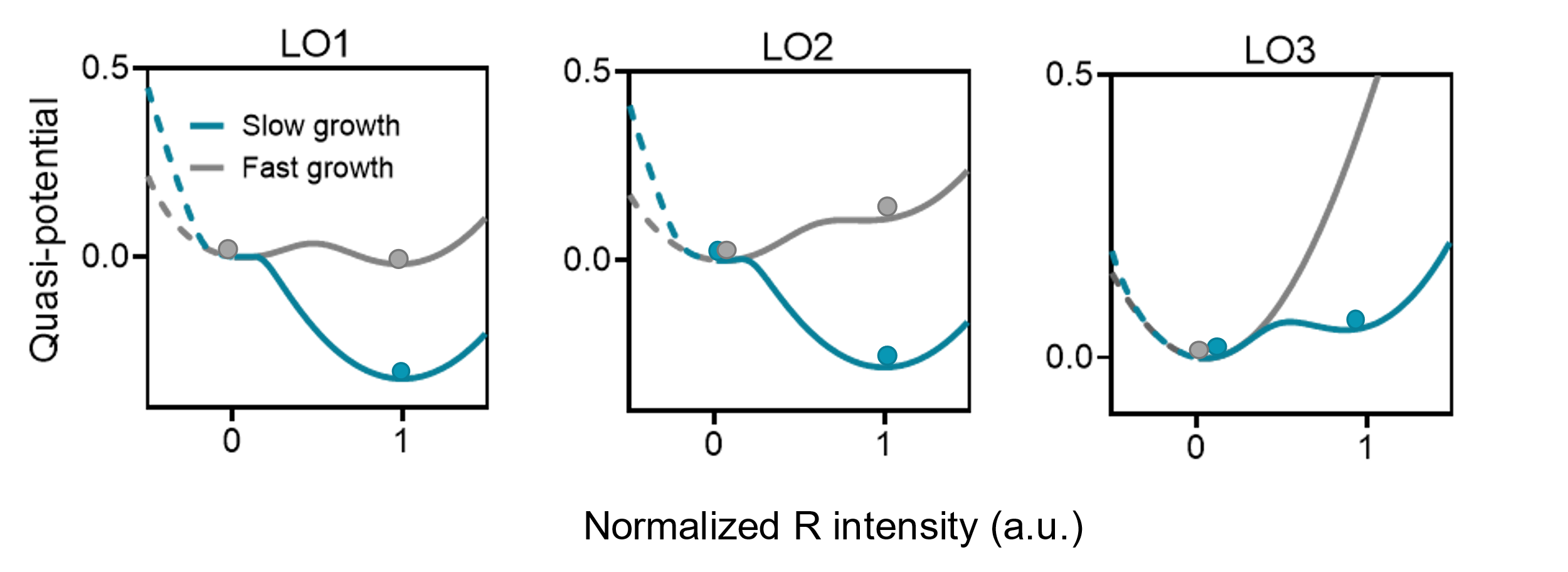

**Supplementary Fig. 3: Quasi-potential landscapes for different strains from Fig. 3b under fast growth (1.60 h^-1^) and slow growth (0.18 h^-1^) conditions.** Circles indicate the basins of attraction (the stable states) under given conditions. It is shown that the LO1 strain displays bistability under fast growth conditions but monostable (R state) under slow growth. LO2 strain features always bistability under growth conditions considered. LO3 strain shows a G state monostable state under fast growth and bistable under slow growth conditions.

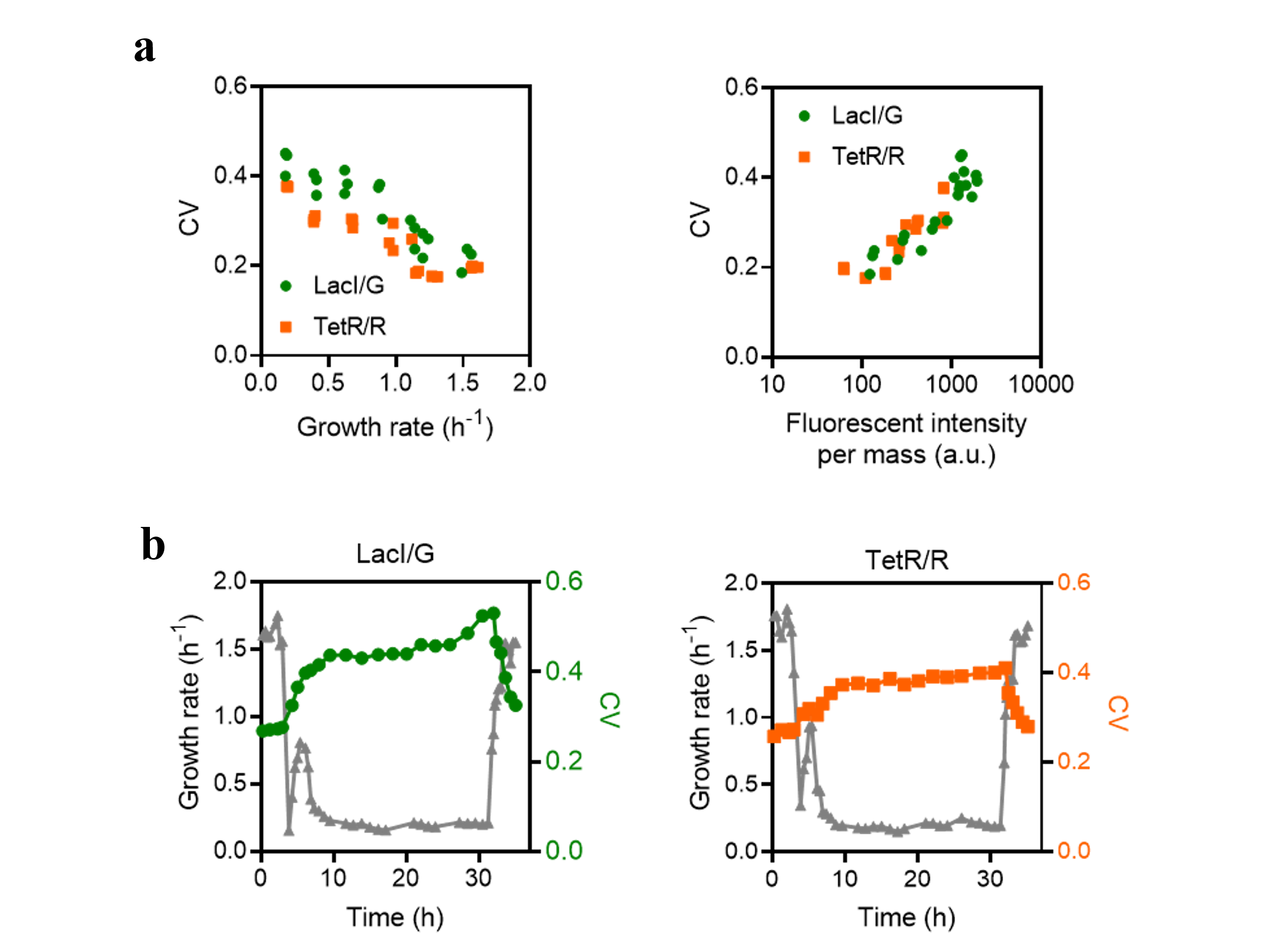

**Supplementary Fig. 4: Gene expression noise is higher at a slower growth rate and higher gene expression level.** Flow cytometry experiments were conducted to measure the CV value (coefficient of variation) of constitutively expressed lacI and tetR (Extended Data Fig. 4a), (**a**) under balanced growth across different conditions, and (**b**) under fluctuating environment (growth downshift and upshift). Similar results were observed by Keren et al.^15^

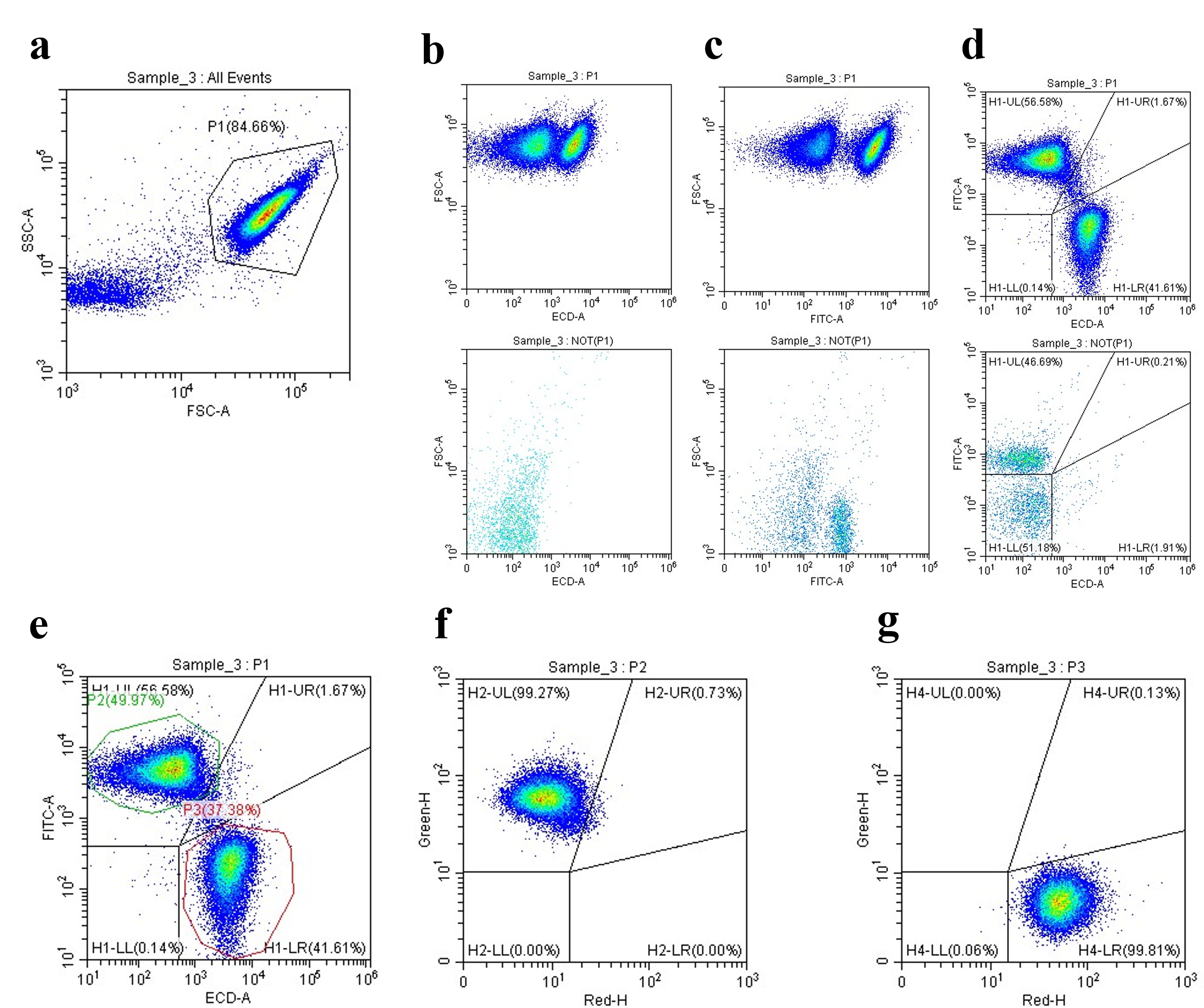

Supplementary Fig 5: Gating strategy used in flow cytometry analysis.

(a) Particles in P1 were regarded as bacterial cells. Cells harbouring the mutual repressive circuit can display a high GFP expression state, high RFP expression, or co-expression state in some cases. An example with a mixture of states is indicated. The FSC-A against FITC-A and FSC-A against ECD-A plots for P1 and NOT(P1) sub-populations are shown in panels b and c, respectively. Panel d is shown the plot of FITC-A/ECD-A, which is divided into subpopulations corresponding to a high GFP expression, a co-expression, a high RFP expression and the non-expression states. Cells in P2 and P3 subpopulations were considered as the green state and the red state, respectively (panel e). (f-g) Plots of GFP intensity per mass against RFP intensity per mass using customized parameters (see Methods) show two distinct states.

**Supplementary Tables**

Supplementary Table 1: Strains used in this study.

| **Strain** | **Parent** | **Genotype** | **Source** |
| --- | --- | --- | --- |
| NCM3722 |  | wild-type *E. coli* K12 strain | Liu Lab |
| NH3 | NCM3722 | NCM3722 Δ*lacIZYA* Δ*fliC* | This study |

Supplementary Table 2: Plasmids constructed in this study.

| **Plasmid** | **Ori** | **Antibiotic** | **Function** | **Source** |
| --- | --- | --- | --- | --- |
| pECJ3 | ColE1 | Kanamycin | Mutual repression circuit | Gift from Dr. James Collins (Addgene, # 75465) |
| pECJ3_LO1 | ColE1 | Kanamycin | Mutual repression circuit | This study (https://benchling.com/s/seq-P3dnzP7p86avljVtkGS4?m=slm-XQYZcUOjf2Xk0KwXjOqU) |
| pECJ3_LO2 | ColE1 | Kanamycin | Mutual repression circuit | This study (https://benchling.com/s/seq-dgxIxFvKk133AKeE6GRt?m=slm-1GbjEGXSWyQyfD6S5vY6) |
| pECJ3_LO3 | ColE1 | Kanamycin | Mutual repression circuit | This study https://benchling.com/s/seq-YTp6y5fiZB5j4ozgLldX?m=slm-or9g5cO1ibcPH55avl3E |
| pECJ3_lacI-sfgfp_delta_tetR | ColE1 | Kanamycin | constitutively expressed *lacI* | This study (https://benchling.com/s/seq-nFerQzJefGBgvrLcsjoW?m=slm-8y9q3aDGUKgFEQc3ULCT) |
| pECJ3_tetR-sfgfp_delta_lacI | ColE1 | Kanamycin | constitutively expressed *tetR* | This study (https://benchling.com/s/seq-pjHVOtXsMsqKHjMj6vbD?m=slm-pxfnkjM3etGe4gtVsKNb) |
| pK4.5_C8_LO1 | ColE1 | Kanamycin | Ptrc_LO1 and mVenus NB reporter | This study (https://benchling.com/s/seq-P3dnzP7p86avljVtkGS4?m=slm-XQYZcUOjf2Xk0KwXjOqU) |
| pK4.5_C8_LO2 | ColE1 | Kanamycin | Ptrc_LO2 and mVenus NB reporter | This study (https://benchling.com/s/seq-K1WUDx9RpQCvSFQKOFz4?m=slm-73Jio6MmKVFvLF0EbwLO) |
| pK4.5_C8_LO3 | ColE1 | Kanamycin | Ptrc_LO3 and mVenus NB reporter | This study (https://benchling.com/s/seq-PKwaB1vRL5d4BTqV7cLn?m=slm-znQqrUSGSHecQbSA6Mr9) |
| pK4-blank | ColE1 | Kanamycin | Blank plasmid for measuring fluorescent background | This study (https://benchling.com/s/seq-h1p9SdiHBgdyUPrywnpT?m=slm-u2xNQhkIXp9xDy2cDt3u) |
| pRS3.R11_lacI | p15A | Spectinomycin | *lacI* expression is regulated by *cymR* system | This study (https://benchling.com/s/seq-vUbJTGgdxSfGtL6GHAon?m=slm-yxE7LfirGfgYzETKJaVV) |
| pRS3.R11_mvenusn | p15A | Spectinomycin | Blank plasmid for measuring fluorescent background | This study (https://benchling.com/s/seq-jphWGtaBoJ64vwjck2K5?m=slm-HFsT9fGhDT5AFE4LIZaY) |
| pRS3.R11_B11_mVenusNB | p15A | Spectinomycin | Blank plasmid for measuring fluorescent background | This study (https://benchling.com/s/seq-xP7fUNXckd2cD4CdaJHW?m=slm-gerqDU0iDXNMiorGdl2A) |

Supplementary Table 3: Chemical components of defined media.

| **Medium Name** | **Buffer** | **Carbon source** | **Other suppl.** | **Note** |
| --- | --- | --- | --- | --- |
| RDM glucose | MOPS | 0.4% (w/v)  glucose | AUCG + EZ |  |
| RDM glycerol | MOPS | 0.4% (v/v)  glycerol | AUCG + EZ |  |
| MOPS CAA glucose | MOPS | 0.4% (w/v)  glucose | 0.2% casamino acids |  |
| MOPS CAA glycerol | MOPS | 0.4% (v/v)  glycerol | 0.2% casamino acids |  |
| MOPS glucose | MOPS | 0.4% (w/v)  glucose | / |  |
| MOPS glycerol | MOPS | 0.4% (v/v)  glycerol | / |  |
| MOPS acetate | MOPS | 60mM  sodium acetate | / |  |
| MOPS glucose glutamate | MOPS  (N-C-) | 0.4% (w/v)  glucose | 10 mM glutamate | Without NH_4_Cl |

Supplementary Table 4: Chemical compositions of supplements for growth media.

| **Supplement name** | **Compound** | **Final conc. (mM)** |
| --- | --- | --- |
| AUCG | Adenine | 0.2 |
|  | Uracil | 0.2 |
|  | Cytosine | 0.2 |
|  | Guanine | 0.2 |
| EZ | Alanine | 0.8 |
|  | Arginine | 5.2 |
|  | Asparagine | 0.4 |
|  | Aspartic acid | 0.4 |
|  | Cysteine | 0.1 |
|  | Glutamic acid | 0.6 |
|  | Glutamine | 0.6 |
|  | Glycine | 0.8 |
|  | Histidine | 0.2 |
|  | Isoleucine | 0.4 |
|  | Leucine | 0.8 |
|  | Lysine | 0.4 |
|  | Methionine | 0.2 |
|  | Phenylalanine | 0.4 |
|  | Proline | 0.4 |
|  | Serine | 10 |
|  | Threonine | 0.4 |
|  | Tryptophane | 0.1 |
|  | Tyrosine | 0.2 |
|  | Valine | 0.6 |
|  | Thiamine | 0.01 |
|  | Calcium pantothenate | 0.01 |
|  | *p*-aminobenzoic acid | 0.01 |
|  | *p*-hydroxybenzoic acid | 0.01 |
|  | 2,3-dihydroxybenzoic acid | 0.01 |

Supplementary Table 5: Experimental data from Fig. 1a and Extended Data Fig. 2.

| **Initial green state** | | | | **Initial red state** | | | |
| --- | --- | --- | --- | --- | --- | --- | --- |
| Time (h) | OD_600_^a^ | RFP intensity per mass (a.u.) | GFP intensity per mass (a.u.) | Time (h) | OD_600_^a^ | RFP intensity per mass (a.u.) | GFP intensity per mass (a.u.) |
| 0.00 | 0.0199 | - - | - - | 0.00 | 0.0236 | - - | - - |
| 0.19 | 0.0282 | 8.9 | 43.4 | 0.18 | 0.0326 | 13.5 | 4.9 |
| 0.37 | 0.041 | 8 | 43.2 | 0.37 | 0.046 | 14.2 | 5.1 |
| 0.55 | 0.0569 | 8 | 39.3 | 0.55 | 0.066 | 13.6 | 4.8 |
| 0.74 | 0.0817 | 7.8 | 40.9 | 0.73 | 0.0936 | 14.1 | 5 |
| 0.92 | 0.1144 | 7.7 | 39.9 | 0.92 | 0.1332 | 14.7 | 5.4 |
| 1.11 | 0.198 | 7.4 | 38.7 | 1.10 | 0.215 | 15.4 | 5.3 |
| 1.31 | 0.262 | 10.6 | 39.6 | 1.31 | 0.293 | 18.3 | 6.2 |
| 1.52 | 0.363 | 11.2 | 43.3 | 1.51 | 0.397 | 19.3 | 6.9 |
| 1.78 | 0.486 | 10.7 | 44.6 | 1.77 | 0.54 | 18.8 | 6.9 |
| 2.03 | 0.602 | 11.8 | 44.7 | 2.02 | 0.749 | 19.3 | 8 |
| 2.33 | 0.769 | 12.3 | 41.5 | 2.31 | 0.828 | 18.6 | 8.8 |
| 2.67 | 0.921 | 12.5 | 35.4 | 2.66 | 1.015 | 16.6 | 9.1 |
| 3.12 | 1.127 | 12.9 | 29.1 | 3.12 | 1.207 | 16.1 | 10.6 |
| 3.74 | 1.319 | 14 | 25.4 | 3.73 | 1.375 | 15.9 | 11 |
| 4.52 | 1.472 | 15 | 23.4 | 4.55 | 1.608 | 16.9 | 11.8 |
| 5.52 | 1.733 | 15 | 22.9 | 5.52 | 1.822 | 17.8 | 12.2 |
| 6.40 | 1.916 | 18.4 | 21.6 | 6.39 | 2.122 | 20.5 | 12.1 |
| 7.23 | 2.07 | 23.2 | 19.9 | 7.22 | 2.304 | 24.4 | 11.8 |
| 8.04 | 2.198 | 31 | 18.9 | 8.03 | 2.507 | 30.5 | 11.7 |
| 8.77 | 2.405 | 42.2 | 18 | 8.75 | 2.73 | 39.4 | 11.3 |
| 9.52 | 2.599 | 55.2 | 17.7 | 9.51 | 2.848 | 49.1 | 11.5 |
| 10.10 | 2.682 | 70.9 | 17 | 10.09 | 2.932 | 61.9 | 11.3 |
| 10.83 | 2.724 | 85.3 | 16.7 | 10.82 | 3.136 | 75.2 | 10.9 |
| 11.37 | 2.975 | 108.2 | 16.3 | 11.36 | 3.218 | 99.3 | 10.8 |
| 11.62 | 2.984 | 132.3 | 16.4 | 11.61 | 3.276 | 120.7 | 10.9 |
| 15 | -.- | 3428.3 | 16.9 | 15 | -.- | 2914.3 | 16.5 |
| 20^a^ | -.- | 43.6 | 8.2 | 20^a^ | -.- | 48.9 | 8.4 |

a: A 10$\times$dilution was achieved when measuring the optical density once OD_600_ was greater than 0.4.

b: Cells from 15h were transferred into fresh SOB medium and successive dilutions were achieved once OD_600_ reached 0.2 for another 5 hours.

Supplementary Table 6: Experimental data from Fig. 1c.

| **Growth medium** | **Cell state** | **Growth rate**  **(h^-1^)** | **RFP intensity per mass (a.u.)** | **GFP intensity per mass (a.u.)** |
| --- | --- | --- | --- | --- |
| RDM glucose | R | 1.591 | 40.1 | 4.7 |
|  | G | 1.606 | 8.0 | 51.4 |
| RDM glycerol | R | 1.234 | 92.8 | 6.6 |
|  | G | 1.212 | 11.4 | 118.4 |
| MOPS CAA glycerol | R | 1.166 | 213.1 | 8.5 |
|  | G | 1.103 | 15.5 | 199.5 |
| MOPS glucose | R | 0.909 | 317.8 | 11.1 |
|  | G | 0.859 | 24.9 | 274.3 |
| MOPS glycerol | R | 0.687 | 803.0 | 14.8 |
|  | G | 0.594 | 37.2 | 567.3 |
| MOPS acetate | R | 0.439 | 1374.8 | 14.7 |
|  | R | 0.439 | 1356.5 | 14.6 |
| MOPS (-NH_4_Cl) glucose glutamate | R | 0.175 | 2207.5 | 19.0 |
|  | R | 0.182 | 2235.0 | 19.5 |

Supplementary Table 7: Experimental data from Fig. 2a.

| **Growth medium** | **Symbol** | **Growth rate (h^-1^)** | **Expression capacity (a.u.)** |
| --- | --- | --- | --- |
| RDM glucose |  | 1.588±0.023 (4) | 21.6±0.9 (4) |
|  |  | 1.525±0.029 (4) | 48.7±5.2 (4) |
| RDM glycerol |  | 1.282±0.019 (4) | 41.1±1.6 (4) |
|  |  | 1.218±0.016 (4) | 101.4±12.9 (4) |
| MOPS CAA glycerol |  | 1.150±0.025 (3) | 69.3±0.6 (3) |
|  |  | 1.127±0.017 (3) | 181.8±13.5 (3) |
| MOPS glucose |  | 0.968±0.014 (4) | 97.7±2.5 (4) |
|  |  | 0.884±0.012 (4) | 309.9±27.9 (4) |
| MOPS glycerol |  | 0.678±0.007 (4) | 173.2±0.4 (4) |
|  |  | 0.624±0.011(3) | 517.8±66.3 (3) |
| MOPS acetate |  | 0.390±0.005 (4) | 362.5±13.9 (4) |
|  |  | 0.406±0.012 (4) | 512.5±38.4 (4) |
| MOPS (-NH_4_Cl) glucose glutamate |  | 0.191±0.002 (3) | 315.5±6.5 (3) |
|  |  | 0.182±0.006 (3) | 589.2±66.6 (3) |

Supplementary Table 8: Experimental data from Fig. 3b. The data of the LO1 strain are given in Supplementary Table 6.

| **Growth medium** | **Strain** | **Cell state** | **Growth rate**  **(h^-1^)** | **RFP intensity per mass (a.u.)** | **GFP intensity per mass (a.u.)** |
| --- | --- | --- | --- | --- | --- |
| RDM glucose | LO2 | R | 1.555 | 51.2 | 5.6 |
|  |  | G | 1.591 | 8.3 | 79.0 |
|  | LO3 | G | 1.605 | 7.9 | 82.9 |
|  |  | G | 1.621 | 7.9 | 83.2 |
| RDM glycerol | LO2 | R | 1.244 | 121.3 | 7.7 |
|  |  | G | 1.218 | 13.3 | 184.9 |
|  | LO3 | G | 1.220 | 11.6 | 167.6 |
|  |  | G | 1.212 | 11.7 | 187.0 |
| MOPS glucose | LO2 | R | 0.917 | 277.6 | 11.2 |
|  |  | G | 0.822 | 24.4 | 507.8 |
|  | LO3 | G | 0.829 | 20.7 | 439.0 |
|  |  | G | 0.811 | 20.8 | 500.9 |
| MOPS glycerol | LO2 | R | 0.680 | 620.3 | 14.1 |
|  |  | G | 0.529 | 39.8 | 1075.4 |
|  | LO3 | G | 0.540 | 29.9 | 869.0 |
|  |  | G | 0.507 | 32.9 | 1150.1 |
| MOPS acetate | LO2 | R | 0.427 | 1705.0 | 13.2 |
|  |  | G | 0.398 | 123.7 | 569.8 |
|  | LO3 | R | 0.432 | 1720.5 | 14.9 |
|  |  | G | 0.379 | 110.1 | 1325.5 |
| MOPS  (-NH_4_Cl) glucose glutamate | LO2 | R | 0.184 | 1336.1 | 15.1 |
|  |  | G | 0.189 | 152.6 | 284.7 |
|  | LO3 | R | 0.186 | 1464.7 | 14.9 |
|  |  | G | 0.189 | 77.6 | 693.4 |

Supplementary Table 9: Parameters used in models.

| **Parameter** | **Description** | **Value** | **Source** |
| --- | --- | --- | --- |
| $\alpha_{R}$ | Growth-rate-dependent protein synthesis rate of TetR-RFP | $\alpha_{R}=\left[ 26.84+\frac{320.22}{\left( 1+\left( \frac{\lambda}{0.66} \right)^{4.09} \right)} \right]\times\lambda$ | Fitted parameters |
| $\alpha_{G}$ | Growth-rate-dependent protein synthesis rate of LacI-GFP | $\alpha_{G}=\left[ 16.61+\frac{627.75}{\left( 1+\left( \frac{\lambda}{0.87} \right)^{4.64} \right)} \right]\times\lambda$ | Fitted parameters |
| $K_{DG,LO1}$ | Dissociation constant of LacI tetramer to its binding site LO1 | 324 | This study, fitted data ^a^ |
| $K_{DG,LO2}$ | Dissociation constant of LacI tetramer to its binding site LO2 | 154.3 | This study, fitted data ^a^ |
| $K_{DG,LO3}$ | Dissociation constant of LacI tetramer to its binding site LO3 | 61.7 | This study, fitted data^a^ |
| $K_{DR}$ | Dissociation constant of TetR dimer to its binding site | 450 | This study, free parameter^a^ |
| $n_{G}$ | Hill coefficient, the cooperativity of repression of promoter Ptrc2 | 4 | Theoretical value^b^ |
| $n_{R}$ | Hill coefficient, the cooperativity of repression of promoter P_L_tetO-1 | 2 | Theoretical value |
| $\tau_{R,LO1}$ | The basal leakage expression level for promoter Ptrc_LO1 | 0.035 | This study, fitted data |
| $\tau_{R,LO2}$ | The basal leakage expression level for promoter Ptrc_LO1 | 0.014 | This study, fitted data |
| $\tau_{R,LO3}$ | The basal leakage expression level for promoter Ptrc_LO1 | 0.007 | This study, fitted data |
| $\tau_{G}$ | The basal leakage expression level for promoter P_L_tetO-1 | 0.002 | This study, free parameter |
| $\kappa_{t}$ | Ribosome elongation rate | $\frac{2.2\cdot\kappa_{R}\cdot\phi_{R}}{\phi_{R}+\phi_{R,0}}$ | Fitted parameters, Data was collected from ref. ^16^ |
| $\phi_{R}$ | Mass fraction of ribosomal proteins | $\lambda/\kappa_{R}+\phi_{R,0}$ | Ref.^1^ |
| $\left[ r_{a} \right]$ | Active ribosome concentration | $\frac{2.2 \cdot\phi_{R}}{1+\left( 0.15/\lambda\right)^{1.36}}$ | Fitted parameters, Data was collected from ref.^16^ |
| $\zeta_{G}$ | Translation efficiency of LacI | 9E3 | This study |
| $\zeta_{R}$ | Translation efficiency of TetR | 5E3 | This study |
| $\phi_{R,0}$ | The vertical intercept of | 0.05 | Ref.^1^ |
| $\kappa_{R}$ | The maximal translation rate of the ribosome | 10.4 | Ref.^1^ |
| $\phi_{G}^{m}$ | mRNA abundance of *lacI* | $0.11\frac{1}{1+\left( \lambda/1.8 \right)^{1.1}}\frac{1}{1+\left( \lambda/0.74 \right)^{3.9}}\cdot$ $\frac{1}{1+\left( 0.71/\lambda\right)^{1.6}}$ | Fitted parameters |
| $\phi_{R}^{m}$ | mRNA abundance of *tetR* | $0.09\frac{1}{1+\left( \lambda/9.5 \right)^{3.3}}\frac{1}{1+\left( \lambda/0.54 \right)^{2.9}}\cdot$ $\frac{1}{1+\left( 0.33/\lambda\right)^{1.6}}$ | Fitted parameters |

a: Dissociation constant: $K_{DR}$ is a free parameter, we fix it at a value of 450. $K_{DG,LO1}$ was the fitted value according to the experimental data (Supplementary Table 6), the magnitudes of $K_{DG,LO2}$ and $K_{DG,LO3}$ were determined according to the experimental relationship (Supplementary Table 8).

b: Instead of using the experimental fitted value 1.15 (Extended Data Fig. 7), we use the theoretical value of 4. The difference between the theoretical value and experimental result may be caused by system errors. Ideally, the fluorescent intensity of LacI-mVenus chimeric protein is linear to the concentration of LacI, however, some factors, such as gene length variation, may affect the plasmid copy number, and mRNA lifetime. These uncertainties can bias the consistency between the fluorescent intensity and the protein concentration. This measuring error results in the bias of hill coefficient.

Supplementary Table 10: Parameters of probability potential landscape.

| **Parameter** | **Description** | **Value** |
| --- | --- | --- |
| $D_{G},D_{R}$ | Magnitudes of noise, the range of growth rate λ is [1.8, 1.5) h^-1^. | 1, 1 |
|  | Magnitudes of noise, the range of growth rate λ is [1.5, 1.0) h^-1^. | 2, 2 |
|  | Magnitudes of noise, the range of growth rate λ is [1.0, 0.6) h^-1^. | 5, 5 |
|  | Magnitudes of noise, the range of growth rate λ is [0.6, 0.1) h^-1^. | 20, 20 |
| $\sigma_{G}^{2}$ | Variance for LacI. | 5^a^ |
| $\sigma_{R}^{2}$ | Variance for TetR. | 5^a^ |

Note: Other parameters that characterize the toggle switch are listed in Supplementary Table 9.

a: the variances of the gaussian distribution do not affect the steady distribution $P\left( G,R,t\to\infty\right)$.

Supplementary Table 11: Functional parts used in this study.

| **Part name** | **Sequence (5’→ 3’)** | **Function** |
| --- | --- | --- |
| PLtetO_M5 | TCCCAATCAGTGATTGAGATTGACATCCCTATCAGTGATAGAGATACTGAGCACATCAGCAGGACGCACTGACCGGATCCATAGGTCC | Promoter |
| Ptrc | TTGACAATTAATCATCCGGCTCGTATAATGTGTGGAATTGTGAGCGGATAACAA | Promoter |
| Ptrc_LO2 | TTGACAATTAATCATCCGGCTCGTATAATGTGTGGAAATGTGAGCGAGTAACAA | Promoter |
| Ptrc_LO1 | TTGACAATTAATCATCCGGCTCGTATAATGTGTGGAATTGTTACTCGCTCACAT | Promoter |
| Ptrc_LO3 | TTGACAATTAATCATCCGGCTCGTATAATGTGTGGAAAATTGTGAGCGCTCACAATT | Promoter |
| *mVenus_NB* | ATGGTTTCTAAAGGTGAAGAACTGTTCACCGGTGTTGTTCCGATCCTGGTTGAACTGGACGGTGACGTTAACGGTCACAAATTCTCTGTTTCTGGTGAAGGTGAAGGTGACGCTACCTACGGTAAACTGACCCTGAAACTGATCTGCACCACCGGTAAACTGCCGGTTCCGTGGCCGACCCTGGTTACCACCCTGGGTTACGGTGTTCAGTGCTTCGCTCGTTACCCGGACCACATGAAACAGCACGACTTCTTCAAATCTGCTATGCCGGAAGGTTACGTTCAGGAACGTACCATCTTCTTCAAAGACGACGGTAACTACAAAACCCGTGCTGAAGTTAAATTCGAAGGTGACACCCTGGTTAACCGTATCGAACTGAAAGGTATCGACTTCAAAGAAGACGGTAACATCCTGGGTCACAAACTGGAATACAACTACAACTCTCACAACGTTTACATCACCGCTGACAAACAGAAAAACGGTATCAAAGCTAACTTCAAAATCCGTCACAACATCGAAGACGGTGGTGTTCAGCTGGCTGACCACTACCAGCAGAACACCCCGATCGGTGACGGTCCGGTTCTGCTGCCGGACAACCACTACCTGTCTTACCAGTCTAAACTGTCTAAAGACCCGAACGAAAAACGTGACCACATGGTTCTGCTGGAATTCGTTACCGCTGCTGGTATCACCCTGGGTATGGACGAACTGTACAAATAA | CDS |
| *gfp_mut2* | ATGCGTAAAGGAGAAGAACTTTTCACTGGAGTTGTCCCAATTCTTGTTGAATTAGATGGTGATGTTAATGGGCACAAATTTTCTGTCAGTGGAGAGGGTGAAGGTGATGCAACATACGGAAAACTTACCCTTAAATTTATTTGCACTACTGGAAAACTACCTGTTCCGTGGCCAACACTTGTCACTACTTTCGGTTATGGTGTTCAATGCTTTGCGAGATACCCAGATCACATGAAACAGCATGACTTTTTCAAGAGTGCCATGCCCGAAGGTTACGTACAGGAAAGAACTATATTTTTCAAAGATGACGGGAACTACAAGACACGTGCTGAAGTCAAGTTTGAAGGTGATACCCTTGTTAATAGAATCGAGTTAAAAGGTATTGATTTTAAAGAAGATGGAAACATTCTTGGACACAAATTGGAATACAACTATAACTCACACAATGTATACATCATGGCAGACAAACAAAAGAATGGAATCAAAGTTAACTTCAAAATTAGACACAACATTGAAGATGGAAGCGTTCAACTAGCAGACCATTATCAACAAAATACTCCGATTGGCGATGGCCCTGTCCTTTTACCAGACAACCATTACCTGTCCACACAATCTGCCCTTTCGAAAGATCCCAACGAAAAGAGAGACCACATGGTCCTTCTTGAGTTTGTAACCGCTGCTGGGATTACACATGGCATGGATGAACTATACAAATAA | CDS |
| *lacI* | ATGGTGAATGTGAAACCAGTAACGCTGTACGATGTCGCAGAGTATGCCGGTGTCTCTTATCAGACCGTTTCCCGCGTGGTGAACCAGGCCAGCCACGTTTCTGCGAAAACGCGGGAAAAAGTGGAAGCGGCGATGGCGGAGCTGAATTACATTCCCAACCGCGTGGCACAACAACTGGCGGGCAAACAGTCGTTGCTGATTGGCGTTGCCACCTCCAGTCTGGCCCTGCACGCGCCGTCGCAAATTGTCGCGGCGATTAAATCTCGCGCCGATCAACTGGGTGCCAGCGTGGTGGTGTCGATGGTAGAACGAAGCGGCGTCGAAGCCTGTAAAGCAGCGGTTCACAATCTTCTCGCGCAGCGCGTCAGTGGGCTGATCATTAACTATCCGCTGGATGACCAGGATGCCATTGCTGTGGAAGCTGCCTGCACTAATGTTCCGGCGTTATTTCTTGATGTCTCTGACCAGACACCCATCAACAGTATTATTTTCTCCCATGAAGACGGTACGCGACTGGGCGTGGAGCATCTGGTCGCATTGGGTCACCAGCAAATCGCGCTGTTAGCGGGCCCATTAAGTTCTGTCTCGGCGCGTCTGCGTCTGGCTGGCTGGCATAAATATCTCACTCGCAATCAAATTCAGCCGATAGCGGAACGGGAAGGCGACTGGAGTGCCATGTCCGGTTTTCAACAAACCATGCAAATGCTGAATGAGGGCATCGTTCCCACTGCGATGCTGGTTGCCAACGATCAGATGGCGCTGGGCGCAATGCGCGCCATTACCGAGTCCGGGCTGCGCGTTGGTGCGGACATCTCGGTAGTGGGATACGACGATACCGAAGACAGCTCATGTTATATCCCGCCGTTAACCACCATCAAACAGGATTTTCGCCTGCTGGGGCAAACCAGCGTGGACCGCTTGCTGCAACTCTCTCAGGGCCAGGCGGTGAAGGGCAATCAACTGTTGCCCGTCTCACTGGTGAAAAGAAAAACCACCCTGGCTCCCAATACGCAAACCGCCTCTCCCCGCGCGTTGGCCGATTCATTAATGCAACTGGCACGACAGGTTTCCCGACTGGAAAGCGGGCAGGCGGCGAACAAAAACGAAGAAAACACCAACGAAGTGCCGACCTTTATGCTGAACGCGGGCCAGGCGAACAGAAGACGAGTTTAA | CDS |
| *tetR* | ATGTCTCGTTTAGATAAAAGTAAAGTGATTAACAGCGCATTAGAGCTGCTTAATGAGGTCGGAATCGAAGGTTTAACAACCCGTAAACTCGCCCAGAAGCTAGGTGTAGAGCAGCCTACATTGTATTGGCATGTAAAAAATAAGCGGGCTTTGCTCGACGCCTTAGCCATTGAGATGTTAGATAGGCACCATACTCACTTTTGCCCTTTAGAAGGGGAAAGCTGGCAAGATTTTTTACGTAATAACGCTAAAAGTTTTAGATGTGCTTTACTAAGTCATCGCGATGGAGCAAAAGTACATTTAGGTACACGGCCTACAGAAAAACAGTATGAAACTCTCGAAAATCAATTAGCCTTTTTATGCCAACAAGGTTTTTCACTAGAGAATGCATTATATGCACTCAGCGCTGTGGGGCATTTTACTTTAGGTTGCGTATTGGAAGATCAAGAGCATCAAGTCGCTAAAGAAGAAAGGGAAACACCTACTACTGATAGTATGCCGCCATTATTACGACAAGCTATCGAATTATTTGATCACCAAGGTGCAGAGCCAGCCTTCTTATTCGGCCTTGAATTGATCATCTGCGGATTAGAAAAACAACTTAAATGTGAAAGTGGGTCTTGA | CDS |
| *mCherry* | ATGGTGAGCAAGGGCGAGGAGGATAACATGGCCATCATCAAGGAGTTCATGCGCTTCAAGGTTCACATGGAGGGCTCCGTGAACGGCCACGAGTTCGAGATCGAGGGCGAGGGCGAGGGCCGCCCCTACGAGGGCACCCAGACCGCCAAGCTGAAGGTGACCAAGGGTGGCCCCCTGCCCTTCGCCTGGGACATCCTGTCCCCTCAGTTCATGTACGGCTCCAAGGCCTACGTGAAGCACCCCGCCGACATCCCCGACTACTTGAAGCTGTCCTTCCCCGAGGGCTTCAAGTGGGAGCGCGTGATGAACTTCGAGGACGGCGGCGTGGTGACCGTGACCCAGGACTCCTCCCTGCAAGACGGCGAGTTCATCTACAAGGTGAAGCTGCGCGGCACCAACTTCCCCTCCGACGGCCCCGTAATGCAGAAGAAGACTATGGGCTGGGAGGCCTCCTCCGAGCGGATGTACCCCGAGGACGGCGCGCTGAAGGGCGAGATCAAGCAGAGGCTGAAGCTGAAGGACGGCGGCCACTACGACGCTGAGGTCAAGACCACCTACAAGGCCAAGAAGCCCGTGCAACTGCCCGGCGCGTACAACGTCAACATCAAGTTGGACATCACCTCCCACAACGAGGACTACACCATCGTGGAACAGTACGAACGCGCCGAGGGCCGCCACTCCACCGGCGGCATGGACGAGCTGTATAAGTAA | CDS |
| *speR* | ATGAGGGAAGCGGTGATCGCCGAAGTATCGACTCAACTATCAGAGGTAGTTGGCGTCATCGAGCGCCATCTCGAACCGACGTTGCTGGCCGTACATTTGTACGGCTCCGCAGTGGATGGCGGCCTGAAGCCACACAGTGATATTGATTTGCTGGTTACGGTGACCGTAAGGCTTGATGAAACAACGCGGCGAGCTTTGATCAACGACCTTTTGGAAACTTCGGCTTCCCCTGGAGAGAGCGAGATTCTCCGCGCTGTAGAAGTCACCATTGTTGTGCACGACGACATCATTCCGTGGCGTTATCCAGCTAAGCGCGAACTGCAATTTGGAGAATGGCAGCGCAATGACATTCTTGCAGGTATCTTCGAGCCAGCCACGATCGACATTGATCTGGCTATCTTGCTGACAAAAGCAAGAGAACATAGCGTTGCCTTGGTAGGTCCAGCGGCGGAGGAACTCTTTGATCCGGTTCCTGAACAGGATCTATTTGAGGCGCTAAATGAAACCTTAACGCTATGGAACTCGCCGCCCGACTGGGCTGGCGATGAGCGAAATGTAGTGCTTACGTTGTCCCGCATTTGGTACAGCGCAGTAACCGGCAAAATCGCGCCGAAGGATGTCGCTGCCGACTGGGCAATGGAGCGCCTGCCGGCCCAGTATCAGCCCGTCATACTTGAAGCTAGACAGGCTTATCTTGGACAAGAAGAAGATCGCTTGGCCTCGCGCGCAGATCAGTTGGAAGAATTTGTCCACTACGTGAAAGGCGAGATCACCAAGGTAGTCGGCAAATAA | CDS |
| *kanR* | ATGATTGAACAAGATGGATTGCACGCAGGTTCTCCGGCGGCTTGGGTGGAGAGGCTATTCGGCTATGACTGGGCACAACAGACAATCGGCTGCTCTGATGCCGCCGTGTTCCGGCTGTCAGCGCAGGGTCGCCCGGTTCTTTTTGTCAAGACCGACCTGTCCGGTGCCCTGAATGAACTGCAAGACGAGGCAGCGCGGCTATCGTGGCTGGCCACGACGGGCGTTCCTTGCGCGGCTGTGCTCGACGTTGTCACTGAAGCGGGAAGGGACTGGCTGCTATTGGGCGAAGTGCCGGGGCAGGATCTCCTGTCATCTCACCTTGCTCCTGCCGAGAAAGTATCCATCATGGCTGATGCAATGCGGCGGCTGCATACGCTTGATCCGGCTACCTGCCCATTCGACCACCAAGCGAAACATCGCATCGAGCGAGCACGTACTCGGATGGAAGCCGGTCTTGTCGATCAGGATGATCTGGACGAAGAGCATCAGGGGCTCGCGCCAGCCGAACTGTTCGCCAGGCTCAAGGCGCGTATGCCCGACGGCGAGGATCTCGTCGTGACCCACGGCGATGCCTGCTTGCCGAATATCATGGTGGAAAATGGCCGCTTTTCTGGATTCATCGACTGTGGCCGGCTGGGTGTGGCGGACCGCTATCAGGACATAGCGTTGGCTACCCGTGATATTGCTGAAGAGCTTGGCGGCGAATGGGCTGACCGCTTCCTCGTGCTTTACGGTATCGCCGCTCCCGATTCGCAGCGCATCGCCTTCTATCGCCTTCTTGACGAGTTCTTCTGA | CDS |
| *cymR_AM* | ATGAGCCCGAAACGTCGTACCCAGGCAGAACGTGCAATGGAAACCCAGGGTAAACTGATTGCAGCAGCACTGGGTGTTCTGCGTGAAAAAGGTTATGCAGGTTTTCGTATTGCAGATGTTCCGGGTGCAGCCGGTGTTAGCCGTGGTGCACAGAGCCATCATTTTCCGACCAAACTGGAACTGCTGCTGGCAACCTTTGAATGGCTGTATGAGCAGATTACCGAACGTAGCCGTGCACGTCTGGCAAAACTGAAACCGGAAGATGATGTTATTCAGCAGATGCTGGATGATGCAGCAGAATTTTTTCTGGATGATGATTTTAGCATCGGCCTGGATCTGATTGTTGCAGCAGATCGTGATCCGGCACTGCGTGAAGGTATTCAGCGTACCGTTGAACGTAATCGTTTTGTTGTTGAAGATATGTGGCTGGGTGTGCTGGTGAGCCGTGGTCTGAGCCGTGATGATGCCGAAGATATTCTGTGGCTGATTTTTAACAGCGTTCGTGGTCTGGTAGTTCGTAGCCTGTGGCAGAAAGATAAAGAACGTTTTGAACGTGTGCGTAATAGCACCCTGGAAATTGCACGTGAACGTTATGCAAAATTCAAACGTTGA | CDS |
| J23100 | GCTAGCACTGTACCTAGGACTGAGCTAGCCGTCAA | Promoter |
| RBS_C8 | TGTCTAATGGCGGTTTTCTAA | Ribosome binding site |
| RiboJ | AGCTGTCACCGGATGTGCTTTCCGGTCTGATGAGTCCGTGAGGACGAAACAGCCTCTACAAATAATTTTGTTTAA | Insulator |
| p15A | TTGAGATCGTTTTGGTCTGCGCGTAATCTCTTGCTCTGAAAACGAAAAAACCGCCTTGCAGGGCGGTTTTTCGAAGGTTCTCTGAGCTACCAACTCTTTGAACCGAGGTAACTGGCTTGGAGGAGCGCAGTCACCAAAACTTGTCCTTTCAGTTTAGCCTTAACCGGCGCATGACTTCAAGACTAACTCCTCTAAATCAATTACCAGTGGCTGCTGCCAGTGGTGCTTTTGCATGTCTTTCCGGGTTGGACTCAAGACGATAGTTACCGGATAAGGCGCAGCGGTCGGACTGAACGGGGGGTTCGTGCATACAGTCCAGCTTGGAGCGAACTGCCTACCCGGAACTGAGTGTCAGGCGTGGAATGAGACAAACGCGGCCATAACAGCGGAATGACACCGGTAAACCGAAAGGCAGGAACAGGAGAGCGCACGAGGGAGCCGCCAGGGGGAAACGCCTGGTATCTTTATAGTCCTGTCGGGTTTCGCCACCACTGATTTGAGCGTCAGATTTCGTGATGCTTGTCAGGGGGGCGGAGCCTATGGAAA | Plasmid replication origin |
| ColE1 | TTGAGATCCTTTTTTTCTGCGCGTAATCTGCTGCTTGCAAACAAAAAAACCACCGCTACCAGCGGTGGTTTGTTTGCCGGATCAAGAGCTACCAACTCTTTTTCCGAAGGTAACTGGCTTCAGCAGAGCGCAGATACCAAATACTGTCCTTCTAGTGTAGCCGTAGTTAGGCCACCACTTCAAGAACTCTGTAGCACCGCCTACATACCTCGCTCTGCTAATCCTGTTACCAGTGGCTGCTGCCAGTGGCGATAAGTCGTGTCTTACCGGGTTGGACTCAAGACGATAGTTACCGGATAAGGCGCAGCGGTCGGGCTGAACGGGGGGTTCGTGCAGACAGCCCAGCTTGGAGCGAACGACCTACACCGAACTGAGATACCTACAGCGTGAGCTATGAGAAAGCGCCACGCTTCCCGAAGGGAGAAAGGCGGACAGGTATCCGGTAAGCGGCAGGGTCGGAACAGGAGAGCGCACGAGGGAGCTTCCAGGGGGAAACGCCTGGTATCTTTATAGTCCTGTCGGGTTTCGCCACCTCTGACTTGAGCGTCGATTTTTGTGATGCTCGTCAGGGGGGCGGAGCCTATGGAAA | Plasmid replication origin |
| ECK120017009 | AGTGAAAAGAAAAAAGGCCGCAGAGCGGCCTTTTTAGTTAGATC | Terminator |
| ECK120033736 | AACGCATGAGAAAGCCCCCGGAAGATCACCTTCCGGGGGCTTTTTTATTGCGC | Terminator |
| PcymRC | AACAAACAGACAATCTGGTCTGTTTGTATTATGGAAAATTTTTCTGTATAATAGATTCAACAAACAGACAATCTGGTCTGTTTGTATTAT | Promoter |

The characters underscored represent the repressors’ operators.

Supplementary References

1	Erickson, D. W. et al. A global resource allocation strategy governs growth transition kinetics of Escherichia coli. *Nature* **551**, 119-123, (2017).

2	Cooper, S. & Helmstetter, C. E. Chromosome replication and the division cycle of Escherichia coli Br. *J. Mol. Biol.* **31**, 519-540, (1968).

3	Paulsson, J., Nordström, K. & Ehrenberg, M. Requirements for rapid plasmid ColE1 copy number adjustments: a mathematical model of inhibition modes and RNA turnover rates. *Plasmid* **39**, 215-234, (1998).

4	Zhang, X. & Bremer, H. Control of the Escherichia coli rrnB P1 promoter strength by ppGpp. *J. Biol. Chem.* **270**, 11181-11189, (1995).

5	Zhu, M., Mori, M., Hwa, T. & Dai, X. Disruption of transcription-translation coordination in Escherichia coli leads to premature transcriptional termination. *Nat. Microbiol.* **4**, 2347-2356, (2019).

6	You, C. et al. Coordination of bacterial proteome with metabolism by cyclic AMP signalling. *Nature* **500**, 301-306, (2013).

7	Klumpp, S. & Hwa, T. Growth-rate-dependent partitioning of RNA polymerases in bacteria. *Proc. Natl. Acad. Sci. USA.* **105**, 20245-20250, (2008).

8	Zhu, M., Mori, M., Hwa, T. & Dai, X. Disruption of transcription-translation coordination in Escherichia coli leads to premature transcriptional termination. *Nat Microbiol* **4**, 2347-2356, (2019).

9	Balakrishnan, R. et al. Principles of gene regulation quantitatively connect DNA to RNA and proteins in bacteria. *bioRxiv*, (2021).

10	Scott, M. et al. Interdependence of cell growth and gene expression: origins and consequences. *Science* **330**, 1099-1102, (2010).

11	Bhattacharya, S., Zhang, Q. & Andersen, M. E. A deterministic map of Waddington's epigenetic landscape for cell fate specification. *BMC Syst. Biol.* **5**, 1-12, (2011).

12	Liao, C., Blanchard, A. E. & Lu, T. An integrative circuit-host modelling framework for predicting synthetic gene network behaviours. *Nat. Microbiol.* **2**, 1658-1666, (2017).

13	Wang, J., Xu, L. & Wang, E. Potential landscape and flux framework of nonequilibrium networks: robustness, dissipation, and coherence of biochemical oscillations. *Proc. Natl. Acad. Sci. USA.* **105**, 12271-12276, (2008).

14	Wang, J., Zhang, K., Xu, L. & Wang, E. Quantifying the Waddington landscape and biological paths for development and differentiation. *Proc. Natl. Acad. Sci. USA* **108**, 8257-8262, (2011).

15	Keren, L. et al. Noise in gene expression is coupled to growth rate. *Genome Research* **25**, 1893-1902, (2015).

16	Dai, X. et al. Reduction of translating ribosomes enables Escherichia coli to maintain elongation rates during slow growth. *Nat. Microbiol.* **2**, 16231, (2016).
